# Comprehensive genome annotation of the model ciliate *Tetrahymena thermophila* by in-depth epigenetic and transcriptomic profiling

**DOI:** 10.1101/2024.01.31.578305

**Authors:** Fei Ye, Xiao Chen, Aili Ju, Yalan Sheng, Lili Duan, Khaled A. S. Al-Rasheid, Naomi A. Stover, Shan Gao

**Affiliations:** MOE Key Laboratory of Evolution & Marine Biodiversity and Institute of Evolution & Marine Biodiversity, Ocean University of China, Qingdao 266003, China; Laboratory for Marine Biology and Biotechnology, Qingdao Marine Science and Technology Center, Qingdao 266237, China; Laboratory of Marine Protozoan Biodiversity & Evolution, Marine College, Shandong University, Weihai 264209, China; Suzhou Research Institute, Shandong University, Suzhou 215123, China; Shum Yiu Foon Shum Bik Chuen Memorial Centre for Cancer and Inflammation Research, School of Chinese Medicine, Hong Kong Baptist University, Hong Kong, SAR, China; Zoology Department, College of Science, King Saud University, Riyadh 11451, Saudi Arabia; Department of Biology, Bradley University, Peoria, Illinois 61625, USA

**Author notes:** Corresponding author (Shan Gao,). These authors contributed equally (Fei Ye, Xiao Chen, Aili Ju).

**Keywords:** *Tetrahymena*, genome annotation, untranslated regions (UTRs), epigenetic information, natural antisense transcript (NATs)

## Abstract

The ciliate *Tetrahymena thermophila* is a well-established unicellular model eukaryote, contributing significantly to foundational biological discoveries. Despite its acknowledged importance, current *Tetrahymena* biology studies face challenges due to gene annotation inaccuracy, particularly the notable absence of untranslated regions (UTRs). To comprehensively annotate the *Tetrahymena* macronuclear genome, we collected extensive transcriptomic data spanning various cell stages. To ascertain transcript orientation and transcription start/end sites, we incorporated data of epigenetic marks displaying enrichment towards the 5’ end of gene bodies, including H3 lysine 4 tri-methylation (H3K4me3), H2A.Z, nucleosomes, and N^6^-methyldeoxyadenine (6mA). Additionally, we integrated Nanopore direct sequencing (DRS), strand-specific RNA-seq, and ATAC-seq data. Using a newly-developed bioinformatic pipeline, coupled with manual curation and experimental validation, our work yielded substantial improvements to the current gene models, including the addition of 2,481 new genes, updates to 6,257 existing genes, and the incorporation of 5,917 alternatively spliced isoforms. Furthermore, novel UTR information was annotated for 26,223 high-confidence genes. Intriguingly, 16% of protein-coding genes were identified to have natural antisense transcripts (NATs) characterized by high diversity in alternative splicing, thus offering insights into understanding transcriptional regulation. Our work will enhance the utility of *Tetrahymena* as a robust genetic toolkit for advancing biological research.

## INTRODUCTION

*Tetrahymena thermophila* (hereafter referred to as *Tetrahymena*) is a well-recognized unicellular model eukaryote and serves as a cornerstone for numerous scientific discoveries [1–11]. Like other ciliates, *Tetrahymena* maintains two functionally distinct nuclei, the diploid micronucleus (MIC) containing five pairs of chromosomes and the polyploid macronucleus (MAC) comprising 181 chromosomes [3, 12–16]. Notably, the MIC remains transcriptionally inactive until the occurrence of sexual reproduction (conjugation), while the MAC is active throughout the vegetative stage to meet the cellular demands [17–19]. The MIC and MAC are generated from the same zygotic nucleus during conjugation [18].

The first assembly of the *Tetrahymena* macronuclear genome was reported in 2006, employing the shotgun sequencing technique [20, 21]. Subsequent versions were published in 2008 and 2014, through the application of next-generation sequencing [22, 23]. Most recently, the contiguity of the MAC genome was substantially improved, ultimately leading to a complete assembly, by using the PacBio Single-Molecule Real-Time (SMRT) sequencing technology [24]. Along with advancements in genome assembly, multiple efforts were dedicated to improving gene annotation, using various datasets from complementary DNA (cDNA) library [20], EST [23], microarray [25], and RNA-seq [26], as well as manual curations. All these genome assembly and gene annotation data were deposited in *Tetrahymena* Genome Database (TGD; Ciliate.org) [22], including two major updates TGD2014 [23] and TGD2021 [24].

However, even the most updated gene model (TGD2021) remained incomplete in several aspects. We found many instances where intron-exon boundary junctions were not accurate, annotated protein-coding genes were fusions of two independent transcription units or needed to be fused with others, or the putative genes were not supported by any RNA-seq reads. In addition, the annotation of untranslated regions (UTRs) was strikingly lacking. Few UTRs were included in the original annotation version when microarray was used for genetic target selection [27]. With the help of Illumina RNA-seq data, the number of genes with 5’ UTRs and/or 3’ UTRs increased to 6,676 [26]; these UTRs data, however, were not integrated into subsequent annotation versions. In TGD2014, only 1,447 genes were annotated with UTRs, representing merely ∼5% of all protein-coding genes [23]. In TGD2021, scarcely any genes were annotated with UTR information [24].

In order to optimize the *Tetrahymena* MAC genome annotation, we generated RNA-seq data from different cell stages, accumulating an ultra-deep sequencing dataset to detect low-expression genes and cell stage-specific genes. Most importantly, we incorporated the distribution information of multiple epigenetic marks, including the histone modification H3K4me3 [5, 28], the histone variant H2A.Z [29], nucleosomes [5, 29], and N^6^-adenine DNA methylation (6mA) [28–30]. All these marks displayed the preferential accumulation towards the 5’ end of the gene body, and were thus helpful in determining the gene orientation and predicting transcription start sites (TSSs). We also integrated Nanopore direct sequencing (DRS) data, strand-specific transcriptomic data, and ATAC-seq data. Based on computational prediction, manual editing, and experimental verification, we have produced a comprehensive annotation of protein-coding genes in the MAC genome of *Tetrahymena*, offering improved precision in intron-exon boundaries, TSS, transcription end sites (TESs), and UTRs. We also performed a preliminary analysis of natural antisense transcripts (NATs), which will help to better resolve the regulation of transcription in *Tetrahymena*.

## RESULTS & DISCUSSION

### Optimize gene model with transcriptomic data

To validate the TGD2021 gene models and identify potential novel genes in the *Tetrahymena* MAC genome, we assembled RNA-seq data from different cell stages, including growth, multiple timepoints during starvation and conjugation (Additional File 2: Table S1). We initially employed LoReAn2 [31], an integrated annotation pipeline, to annotate protein-coding genes. The average lengths of predicted coding regions (3,900 bp vs. 2,452 bp) and intergenic regions (5,550 bp vs. 1,456 bp) were both considerably longer than those in TGD2021 (Additional File 3: Table S2). However, the number of protein-coding genes was notably lower (15,355 vs. 26,259) and only 8,351 of these genes contained UTR information. Moreover, these predicted coding regions covered only 37.61% of the entire genome (38.87 Mb out of 103.34 Mb), much lower than the coverage in TGD2021 (64.38 Mb, 62.30%) [24]. The incompleteness of the gene annotation was further manifested by the lower mapping ratio of RNA-seq reads (56.74% vs. 82.07% in TGD2021), suggesting that a large proportion of genes were not annotated by LoReAn2. Collectively, the unsuccessful annotation by LoReAn2 [31] prompted us to develop a more efficient approach for the *de novo* annotation of the *Tetrahymena* MAC genome.

Here, we employed a newly developed pipeline (Fig. 1, Additional File 1: Fig. S1), which enabled us to identify a total of 27,369 gene candidates (draft v1). Many gene candidates (17,170 out of 27,369, 63%) shared identical intron-exon boundary junctions with TGD2021 and were thus temporarily considered well-annotated genes. For the remaining gene candidates, we further optimized their annotations using full-length transcripts obtained from Nanopore direct RNA sequencing (DRS), strand-specific RNA-seq (ssRNA-seq), and the most highly expressed RNA-seq transcripts among all cell stages. 3,408 new genes were identified, mostly located within intergenic regions as previously defined in TGD2021 (Fig. 2A, Additional File 1: Fig. S2A).

**Figure 1.**
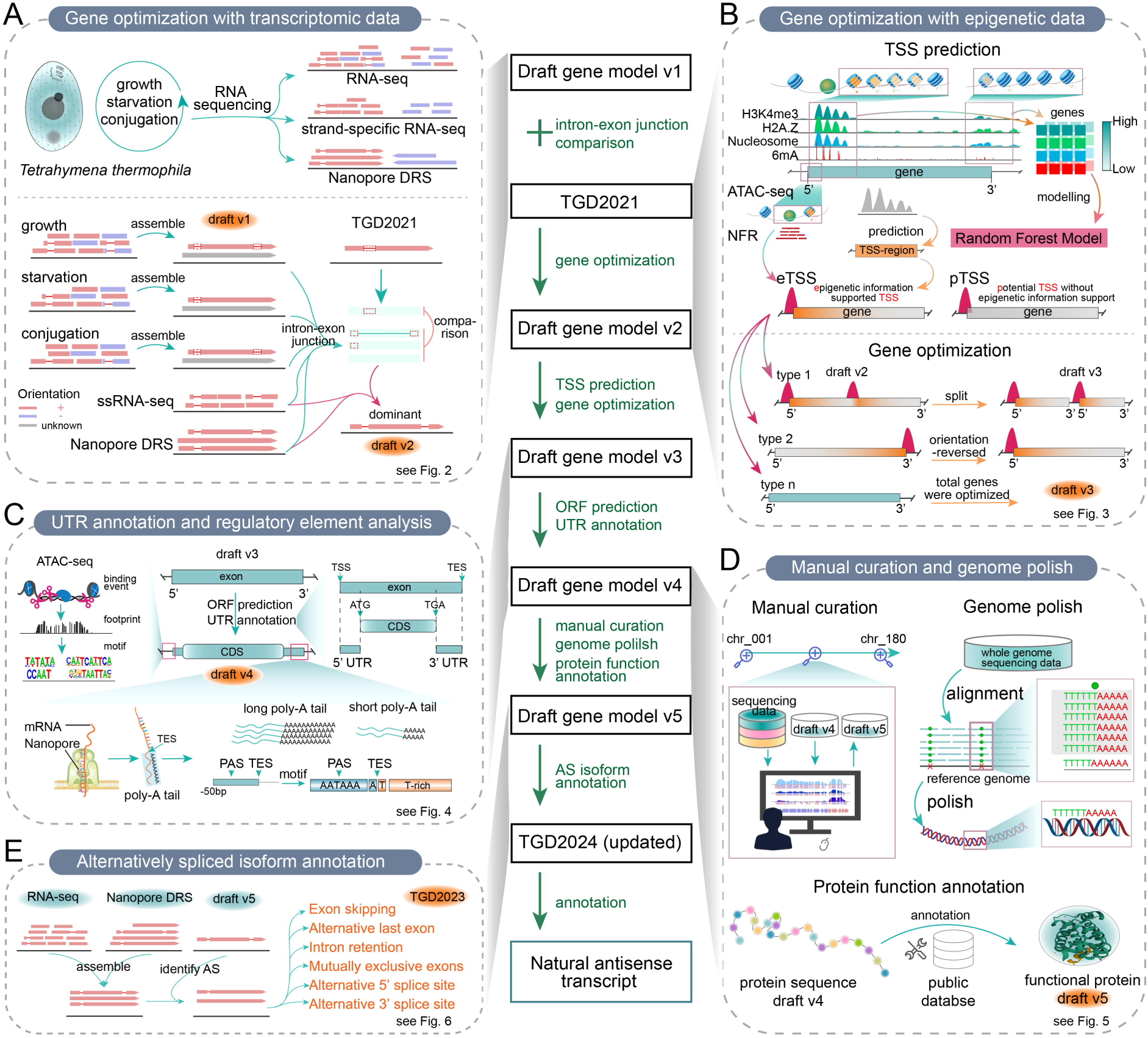
Schematic overview of gene model optimization by integrating transcriptomic and epigenetic data. A. Transcripts at different stages of growth, starvation, and conjugation were assembled into draft v1. By comparing newly assembled transcripts with those from TGD2021, well-annotated genes were retained, and error genes were optimized with the assistance of Nanopore DRS and strand-specific RNA-seq (ssRNA-seq) data, resulting in draft v2. B. Epigenetic data were integrated to predict transcription start sites (TSSs) using a random forest (RF) model, and TSSs were further categorized into eTSSs and pTSSs with the addition of ATAC-seq data. Further optimization of the gene model was achieved using eTSSs, resulting in draft v3. C. Transcripts in draft v3 were subjected to open reading frame (ORF) prediction, and UTR information was provided based on information of CDS, TSSs and transcription end sites (TESs), resulting in draft v4. Features of regulatory elements including promoters, poly-A sequences, and poly-A signals were analyzed. D. The draft gene model v4 underwent two rounds of manual curation, followed by additional genome polish and protein function annotation, resulting in the generation of an improved gene model, draft v5. E. Annotation of alternatively spliced (AS) isoforms was performed by integrating RNA-seq and Nanopore DRS data, resulting in TGD2024 (updated). Natural antisense transcripts (NATs) were annotated based on the updated gene model. TGD: *Tetrahymena* genome database; NFR: nucleosome free region; PAS: poly-A signal.

**Figure 2.**
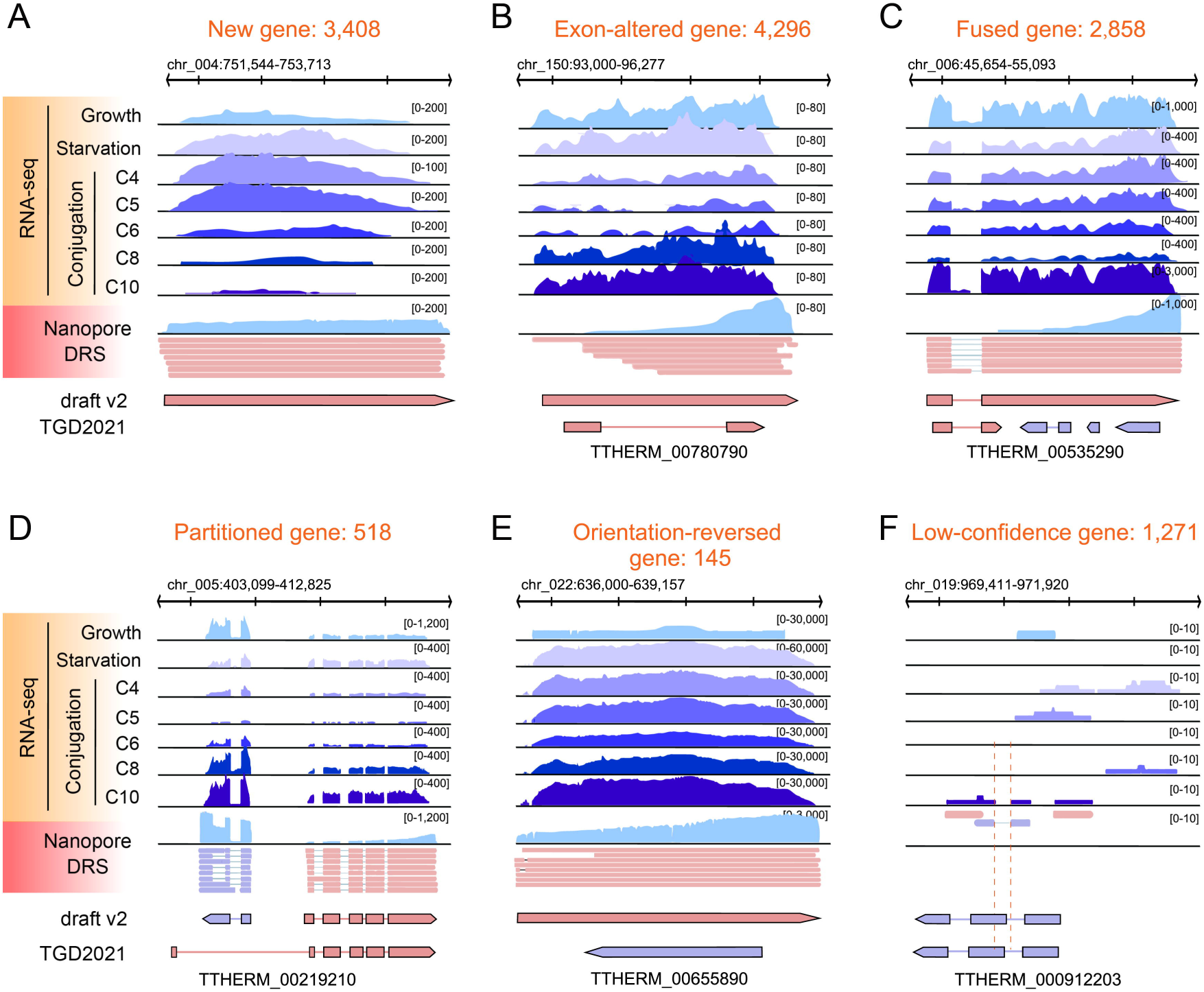
IGV snapshots showing five categories of gene models optimized by transcriptomic data, including new gene (A), exon-altered gene (B), fused gene (C), partitioned gene (D), and orientation-reversed gene (E). Low-confidence genes (F) were not supported by RNA-seq data, thus retaining their annotations in draft v2 as in TGD2021. Tracks from top to bottom were RNA-seq (growth, starvation 24h, and conjugation at 4h, 5h, 6h, 8h, and 10h), Nanopore DRS coverage and reads alignment, and the gene models of draft v2 and TGD2021. Reads and gene models in pink represented transcription on the sense strand, and those in purple on the antisense strand.

Most importantly, we optimized gene annotations for a substantial number of genes (7,817). These optimizations fell into four major classes. 1) Exon-altered genes: 4,296 genes had altered intron-exon boundaries (Fig. 2B, Additional File 1: Fig. S2B). 2) Fused genes: 2,858 genes were merged accordingly and annotated as 1,314 genes. These mergers were supported by RNA-seq reads and DRS reads spanning two neighboring genes (Fig. 2C, Additional File 1: Fig. S2C). 3) Partitioned genes: 518 genes were separated into 1,036 genes, based on RNA-seq reads that were interrupted in the middle of these genes, with no RNA-seq reads spanning the neighboring genes (Fig. 2D, Additional File 1: Fig. S2D). 4) Orientation-reversed genes: the orientation of 145 single-exon genes was changed according to the strand-specific reads (Fig. 2E).

1,271 genes in TGD2021 were defined as low-confidence genes (Fig. 2F). First, there were 86 genes for which no RNA-seq reads were detected among all our data, suggesting that these genes either have extremely low expression levels or their expression is highly specific to certain conditions. The prediction of these genes was more likely attributable to errors in the *ab initio* annotation, given the absence of supports from previous datasets, including EST, microarray, and Illumina RNA-seq data [20, 23, 25, 26]. Second, for 1,185 genes, their sequenced reads did not align well with the original annotation, especially at intron-exon boundaries. Despite the presence of RNA seq reads, new transcripts failed to be assembled owing to its low sequencing depth and occasionally mixed antisense reads.

At this stage, we identified a total of 28,640 genes (draft v2) (Additional File 1: Fig. S1), encompassing 17,170 well-annotated genes, 3,408 new genes, 4,296 exon-altered genes, 1,314 fused genes, 1,036 partitioned genes, 145 orientation-reversed genes, and 1,271 low-confidence genes.

### Refine gene model using epigenetic marks

To further enhance the accuracy of gene models optimized by transcriptomic data, we developed a machine learning algorithm to utilize information from epigenetic marks (Fig. 1B, 3A). These marks, including H3K4me3, H2A.Z, 6mA, and nucleosome positioning, all exhibited the preferential enrichment at the 5’ end of actively transcribed genes [5, 28–30, 32], thus providing valuable guidance for predicting TSSs. For model training and evaluation, 10,460 long genes (> 1kb) were selected from a pool of 17,170 well-annotated genes (see more details in Methods and Materials). Using the trained Random Forest (RF) model (Fig. 3B), 27,840 TSS regions were predicted.

**Figure 3.**
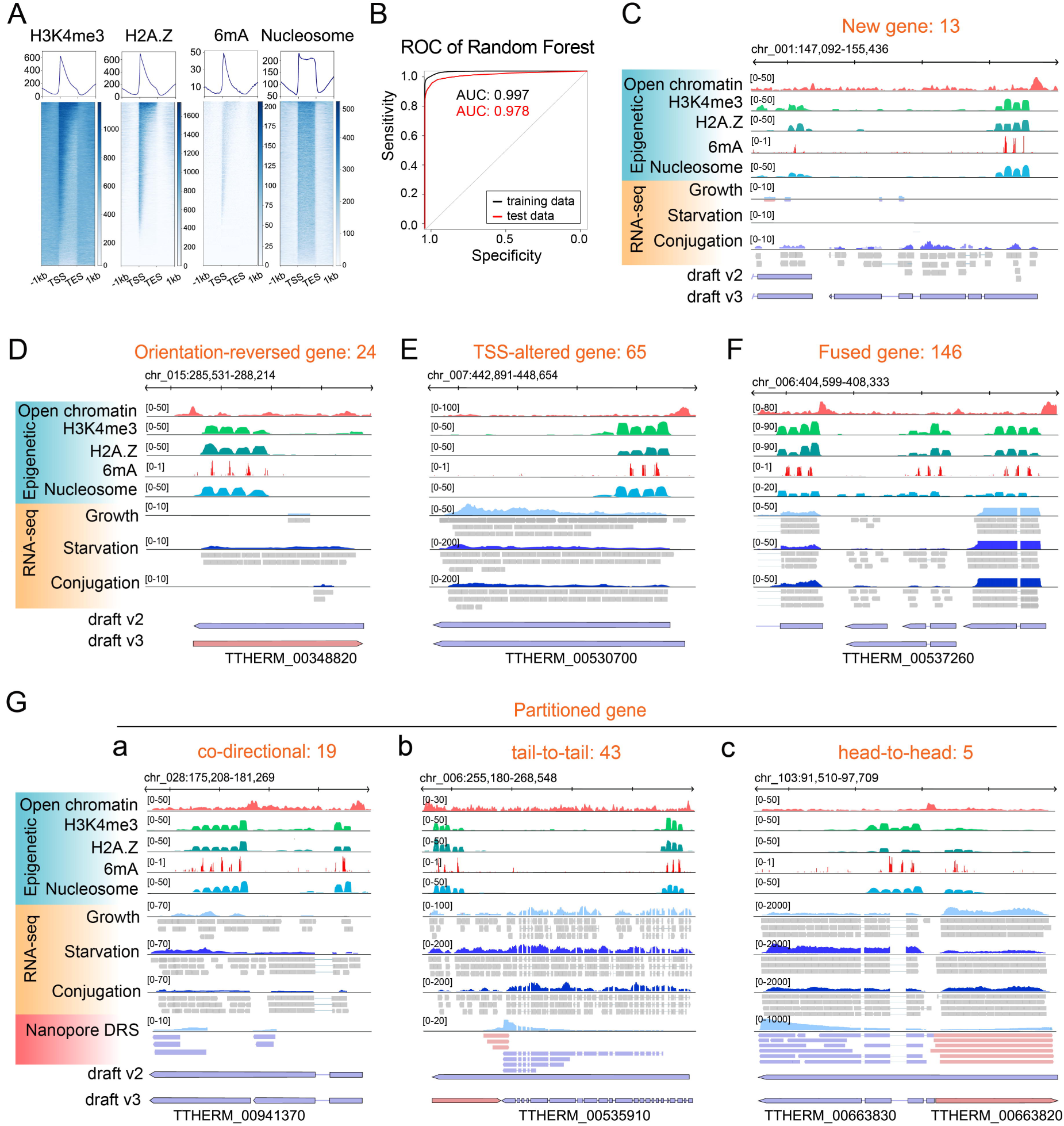
Gene model optimization using epigenetic information. A. Distribution profiles of H3K4me3, H2A.Z, 6mA, and nucleosome on the gene body. Genes were scaled to unit length and was extended to each side by 1kb length. Note that all four marks were accumulated downstream of TSS, towards the 5’ end of the gene body. B. The ROC-AUC curve (ROC: Receiver Operating Characteristics, AUC: Area Under the Curve) measuring the performance of our random forest (RF) model. The ROC was a probability curve and AUC represented the degree or measure of separability. The higher the AUC, the better the model was at predicting “TSS-region” classes as “TSS-region” classes or “not-TSS-region” classes. The AUC for both the training data and the testing data was close to 1, indicating excellent performance of our RF model in predicting TSS-region. C-G. IGV snapshots of seven types of gene models optimized by epigenetic data with the complementation of transcriptomic data, including new gene (C), orientation-reversed gene (D), TSS-altered gene (E), fused gene (F), and partitioned gene (G). Partitioned gene was further subcategorized as co-directional (a), tail-to-tail (b), and head-to-head (c). The tracks from top to bottom were epigenetic information including nucleosome free region (NFR) deduced from ATAC-seq, H3K4me3, H2A.Z, 6mA, and nucleosome, and RNA-seq transcripts of different cell stages. The most highly expressed transcripts in conjugation were selected. Reads and gene models in pink represented transcription on the sense strand, and those in purple on the antisense strand.

ATAC-seq fragments from the nucleosome-free region (NFR) tended to enrich on gene promoters around TSS (Additional File 1: Fig. S3A) [33]. From our ATAC-seq data, we identified 42,469 significant broad peaks at NFR, and the center of each peak was defined as a candidate TSS. Of these candidate TSSs, those located within 200 bp of the TSS regions predicted by our RF model were defined as epigenetic data supported TSSs (eTSSs). Those located within 200 bp of 5’ end of genes, lacking support from our RF model, were defined as potential TSSs (pTSSs) (Additional File 1: Fig. S3B).

Among 28,640 genes optimized by transcriptomic data (draft v2), 25,617 possessed either eTSS (21,095) or pTSS (4,522) (Additional File 1: Fig. S1). 3,937 genes had multiple eTSSs and they were subsequently subjected to manual curation (Fig. 5A, B). Interestingly, 885 head-to-head gene pairs (1,670 genes) shared their respective eTSSs, indicating that they utilized a bidirectional promoter for transcription (Additional File 1: Fig. S3C). We also found 1,752 genes with neither eTSS nor pTSS. These genes were either tandem duplicate genes (Additional File 1: Fig. S3D) or duplicated genes with multiple short exons (Additional File 1: Fig. S3E). These duplicated genes were also subsequently subjected to manual curation (Fig. 5C, D). For the 1,271 low-confidence genes poorly supported by RNA-seq reads, neither eTSS nor pTSS were found in close proximity to them (Additional File 1: Fig. S3F), further confirming that they are either silent genes or error genes [20].

**Figure 4.**
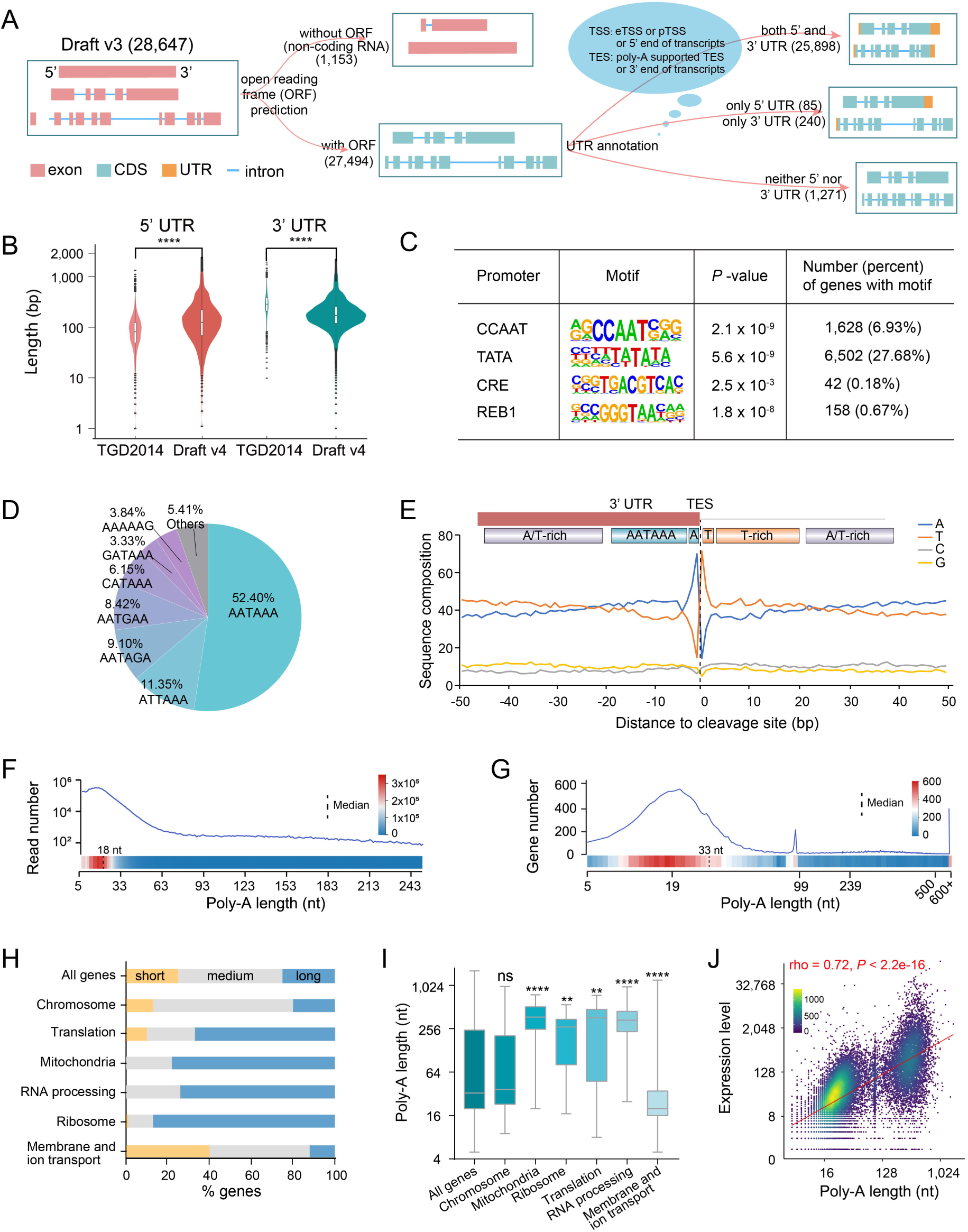
UTR annotation and regulatory elements analysis. A. Schematics for UTR annotation. ORF prediction was conducted on top of draft v3, resulting in a total of 27,494 protein-coding genes. 1,153 genes lacking ORF were defined as potential non-coding RNA. A putative ORF was defined as amino acids sequence longer than 100 aa. UTR information was further supplemented based on predicted TSSs and TESs. 1,271 low-confidence genes defined in Figure 2F lacked UTR annotations. Draft v3 after ORF prediction and UTR annotation generated draft v4. B. UTR comparisons between draft v4 and TGD2014, the latter of which contained UTR information for 1,477 genes. Student’s *t*-test was performed. **** *P*<0.0001. C. Enriched core promoter motifs in promoter proximal sequences around TSS—were identified by Homer [90]. P-values represented the statistical significance of motif enrichment, indicating the likelihood that the observed frequency of each motif in the specified genomic region was greater than what would be expected by chance. D. Venn diagram showing the composition of sequence motifs around poly-A signals (PAS). AATAAA was identified as the most predominant motif. E. Summary of nucleotide frequencies and main regulatory elements around cleavage sites. Cleavage sites were significantly associated with the AT motif. The dashed black line represented positions of cleavage sites. F. Length distribution of poly-A tails identified in Nanopore DRS (minimum reads >5). The median length was 18nt, illustrated by a dashed black line. G. Distribution of the maximal poly-A tail length for each gene (number of gene with poly-A = 21,660). All genes were sorted by the length of their longest poly-A tails from shortest to longest and divided into three groups: 1) the first 25% of genes, defined as short-tailed genes, with poly-A tail length ranging from 5-19 nt; 2) the middle 25%-75% of genes, defined as medium-tailed genes, with poly-A tail length between 19-239 nt; 3) the remaining 25% of genes, defined as long-tailed genes, with poly-A tail length exceeding 239 nt. H. Gene ontology (GO) analysis revealed that short-tailed and long-tailed genes were enriched in distinct functional groups. The colored bars represented the percentage of genes in each tail-length category. I. Distribution of poly-A tail length in different functional groups. Student’s *t*-test was performed. ** *P* <0.01, *** *P* <0.001, and ns *P* >0.05. J. The Spearman’s correlation between poly-A tail length and gene expression level (rho=0.72, *P* <2.2e-16). The longest poly-A tail was selected as the representative for each gene. Gene expression levels were quantified using the number of Nanopore DRS reads, with the removal of interference from antisense RNA. Both axes were plotted on a logarithmic scale.

**Figure 5.**
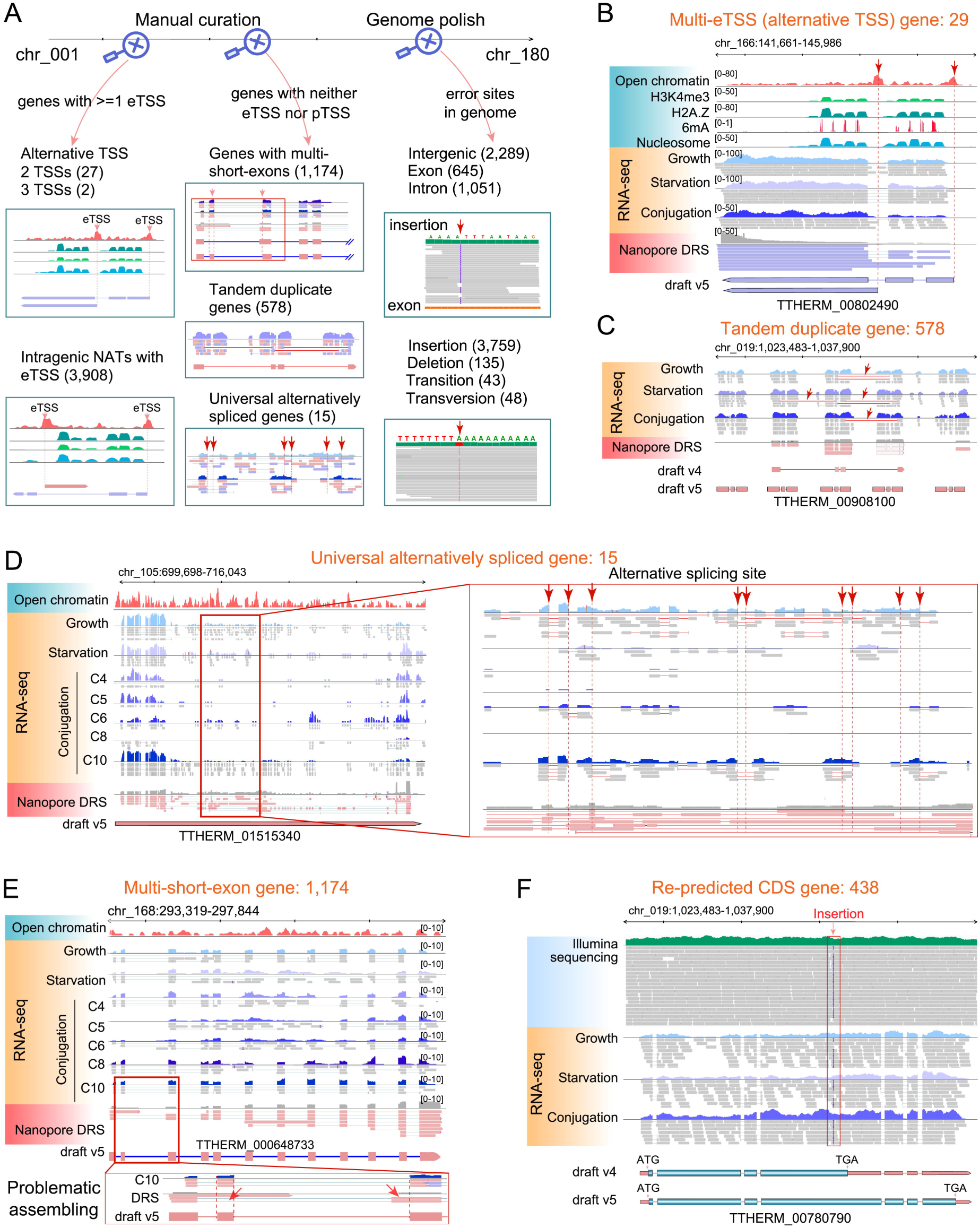
Manual curation, genome polish, and protein function annotation. A. Illustration of manual curation and genome polish on draft v4, resulting in draft v5. Two rounds of manual curation were conducted for all 180 non-rDNA chromosomes, focusing on genes with more than one eTSS, as well as those with neither eTSS nor pTSS. Genome polish was conducted by correcting error sites identified through manual curation using Illumina sequencing data. B. An IGV snapshot showing the manual curation of a multi-eTSS gene, based on epigenetic and transcriptomic data. The tracks from top to bottom were nucleosome free region (NFR), H3K4me3, H2A.Z, 6mA, nucleosome, RNA-seq transcripts from different cell stages, and Nanopore DRS transcripts. The arrows and dashed lines indicated positions of eTSSs. C. An IGV snapshot showing the manual curation of a tandem duplicate gene, by incorporating RNA-seq transcripts of different cell stages with its corresponding reads alignment and Nanopore DRS reads. The arrows indicated the chimeric alignment of RNA-seq transcripts. D. An IGV snapshot showing the manual curation of a universally alternatively spliced gene, by incorporating RNA-seq transcripts of different cell stages with its corresponding reads alignment and Nanopore DRS reads. In the magnified box on the right, arrows indicated the universal alternatively spliced site. These universally alternatively spliced genes were annotated with their most dominant isoforms. E. An IGV snapshot showing that multi-short-exon genes were always error-assembled when using Nanopore DRS data. This manual curation was performed with the aid of RNA-seq data from multiple stages. In the magnified box on the bottom, arrows indicated error-assembled sites. F. An IGV snapshot showing an insertion site located in the exon resulted in erroneous CDS predictions. This manual curation was supported by both Illumina and transcriptomic data. The arrow and box indicated the insertion site.

Based on eTSSs with high confidence, we reexamined the gene model (draft v2) that has been refined by transcriptomic data (Fig. 1B). 13 genes were identified as new genes based on the presence of eTSSs (Fig. 3C, Additional File 1: Fig. S4A). These genes were lowly expressed, limited to only one developmental stage, and were not originally annotated by our pipeline due to the scarcity of supporting reads in the stage-combined RNA-seq dataset.

Meanwhile, annotations of multiple genes were optimized based on eTSS, complemented by transcriptomic data (Fig. 1B). 1) Orientation-reversed genes: the orientation of 24 single-exon genes was reversed, because their eTSSs were located within the previously annotated 3’ UTRs (Fig. 3D). Their orientation could not be determined by ssRNA-seq reads, because they were not expressed during the growth stage when the ssRNA-seq was conducted. 2) TSS-altered genes: the TSSs of 65 genes were altered according to the positions of their eTSSs. The gaps between eTSSs and TSSs predicted by transcriptomic data were attributed to the limited RNA-seq read coverage. Consequently, their TSSs were extended to align with eTSSs, supported by limited yet existing RNA-seq reads (Fig. 3E). 3) Fused genes: 146 genes were merged into 73 genes. These genes were initially misclassified into two separate genes primarily attributed to minor gaps between two clusters of RNA-seq reads. However, only one of the constituent genes contained a well-defined eTSS, while the other lacked any discernible eTSS or pTSS. The surrounding genes each had respective eTSSs, thus eliminating their chances to be merged with other genes (Fig. 3F, Additional File 1: Fig. S4B). 4) Partitioned genes: 67 genes were split into 134 genes. These genes contained two different eTSSs that were divided into three subgroups: (a) co-directional, 19 genes had two eTSSs either simultaneously at the 5’ end and the middle of the previously annotated genes or at the 3’ end and the middle (Fig. 3G, Additional File 1: Fig. S4C), representing two genes transcribed in the same direction; (b) tail-to-tail, 43 genes had two peaks at both 5’ and 3’ ends, respectively (Fig. 3G, Additional File 1: Fig. S4D), representing two convergent genes proceeding in opposite directions and towards each other; and (c) head-to-head, 5 genes had two close yet separated peaks in the middle of the gene body (Fig. 3G, Additional File 1: Fig. S4E), representing two divergent genes proceeding in opposite directions and away from each other.

Compared to draft v2, draft v3 (Additional File 1: Fig. S1) contained 13 new genes and 255 optimized genes including 24 TSS-altered genes, 73 fused genes, 134 partitioned genes, and 24 orientation-reversed genes.

### UTR annotation and regulatory elements analysis

By employing Nanopore DRS data (Additional File 1: Fig. S5A), we identified TES, defined as the 3’ cleavage/polyadenylation site before the poly-A tail [34] [35, 36], in 75% (21,660 out of 28,647) genes. 1,915 genes harbored multiple TESs (Additional File 1: Fig. S5B). For the genes in draft v3 with well-defined TSS and TES, we predicted coding DNA sequences (CDSs) and open reading frames (ORFs) according to the ciliate genetic code [37] for 28,647 genes. 1,153 genes with no predictable ORF were classified as potential non-coding RNA (Fig. 4A). We then defined the regions on both sides of transcripts, excluding the CDS, as 5’ UTRs and 3’ UTRs respectively (Fig. 1C, Fig. 4A). In total, 25,898 genes had both 5’ UTRs and 3’ UTRs, 85 genes had only 5’ UTRs, and 240 genes only had 3’ UTRs. The 1,271 low-confidence genes did not have annotated UTR information (Fig. 4A). The average lengths of 5’ UTRs and 3’ UTRs were 192.54 bp and 238.61 bp (Fig. 4B), respectively. Moreover, the inclusion of more precise and reliable UTR information also increased the mapping ratio of RNA-seq reads (82.07% in TGD2021 vs. 91.87% in the updated gene model).

In the proximal promoter sequences surrounding TSS, we identified several core promoter motifs that may play a role as *cis*-elements (Fig. 4C, Additional File 5: Table S4). They contained key motifs involved in transcription activation, such as CCAAT box [38], TATA box [39], and CRE (cAMP response element) (TGACGTCA) [40], and involved in nucleosome positioning (Reb1: CGGGTAA) [41, 42].

In metazoans, a predominant polyadenylation signal (PAS) was observed within the region spanning 0 and 50 bp upstream of the RNA cleavage site [43–45]. In *Tetrahymena*, PAS also consisted of a primary dominant AATAAA motif, along with six variant motifs ATTAAA, AATGAA, AATAGA, CATAAA, GATAAA, and AAAAAG (Fig. 4D) [43]. However, there was a pronounced AT motif in *Tetrahymena* (Fig. 4E), in contrast to the CA motif at the cleavage site in mammals [46]. In metazoans, GT-rich elements (GTGT) were observed both upstream and downstream of the cleavage site [47]. In *Tetrahymena*, however, T-rich sequences were observed within 20 bp downstream and AT-rich beyond 30 bp upstream (Fig. 4E, Additional File 1: Fig. S5C). This suggests that *Tetrahymena* may have distinct mRNA cleavage and polyadenylation mechanisms compared to metazoans.

Additionally, we found that the length of poly-A tails peaked at approximately 18 nt in *Tetrahymena*, similar to *Arabidopsis* (∼19 nt), soybean (∼19 nt), maize (∼18 nt) and rice (∼18 nt) [48] (Fig. 4F). When analyzing the longest poly-A sequences of each gene, it was observed that their poly-A length of genes exhibited two prominent peaks at 13-30 nt and 95-100 nt, respectively (Fig. 4G). To investigate whether functional classes of genes are associated with the length of poly-A tails, we sorted all genes by the length of their longest poly-A tails from shortest to longest and divided them into three groups: 1) the first 25% of genes, defined as short-tailed genes, with poly-A lengths ranging from 5-19 nt; 2) the middle 25%-75% of genes, defined as medium-tailed genes, with poly-A lengths between 19-239 nt; and 3) the remaining 25% of genes, defined as long-tailed genes, with poly-A lengths exceeding 239 nt (Fig. 4H). Gene ontology (GO) analysis revealed that short-tailed genes were highly enriched in the pathway of membrane and ion transport, while long-tailed genes were more prominently enriched in functions related to mitochondria, translation, RNA processing, and ribosome (Fig. 4H, I, Additional File 6: Table S5). This was in strong contrast to *Caenorhabditis elegans* and mammals, wherein short-tailed genes were highly enriched for genes involved in translation, nucleosome, and ribosome [49]. Additionally, we identified a positive correlation between lengths of gene poly-A tails and their expression levels (rho =0.72, *P* <2.2e-16) (Fig. 4J) in *Tetrahymena*, suggesting that long poly-A tails stabilize mRNA [50, 51]. This contrasted with the previous finding in *Caenorhabditis elegans*, where highly expressed mRNAs were observed to have shorter poly-A tails, explained by enhanced translation efficiency and the maintenance of an optimal tail length [49]. No correlation was observed between gene poly-A length and gene length (rho= -0.089, *P* <2.2e-16) (Additional File 1: Fig. S5D). The discrepancy between *Tetrahymena* and other eukaryotes suggested functional diversification of poly-A tails across different species.

In this version of gene models (draft v4) (Additional File 1: Fig. S1), 25,898 genes had both 5’ UTRs and 3’ UTRs, 85 genes had only 5’ UTRs, 240 genes only had 3’ UTRs, and 1,271 genes had no UTR.

### Manual curation, genome polish, and protein annotation

Subsequently, we performed manual curation in IGV-sRNA [52], conducting two rounds of evaluations across the 180 non-rDNA chromosomes (Fig. 1D, 5A). Firstly, we checked the 3,937 genes with multiple eTSSs. Among the 3,935 genes with two eTSSs, 3,908 were capable of transcribing antisense transcripts, with one eTSS belonging to a protein-coding gene and the other eTSS corresponding to an antisense transcript (Fig. 5A, 7D-F). The remaining 27 genes contained two eTSSs, with one of them serving as an alternative TSS for the protein-coding gene (Fig. 5A, B). Additionally, two genes contained three eTSSs, signifying three alternative TSSs. Secondly, we checked 1,752 duplicated genes with neither eTSS nor pTSS (Fig. 4A). These genes could be categorized into two groups. One group consisted of 1,174 repetitive genes with multiple short exons (mostly <100 bp) distributed across distinct chromosomes (Fig. 5A, 5E). They tended to be misaligned due to the default Smith-Waterman algorithm for Nanopore DRS data analysis [53]. Most of these multi-short-exon genes belonged to the leucine-rich repeat (LRR) superfamily, which has recently evolved and lacks the transcription activation marks including 6mA [54]. The other group comprised 578 tandem duplicate genes with multiple copies arranged in a linear fashion at a single genomic locus (Fig. 5A, C). Thirdly, there were 15 genes exhibiting super high splicing diversity, with nearly every noncoding exon being subject to alternatively splicing (near-universal AS) (Fig. 5D). This phenomenon was also observed in humans, wherein 69% of human protein-coding exons were classified as alternative, and some functional long-noncoding RNAs (lncRNAs) such as XIST, HOTAIR, GOMAFU, and H19 were observed to be near-universally alternatively spliced at each locus [55]. We annotated these 15 genes with their most dominant isoforms.

While conducting manual curation, we observed sequence errors in certain regions. Therefore, we polished the genome sequence using Illumina sequencing data (Figure 1D, 5A), correcting a total of 3,759 insertions, 135 deletions, 43 transitions, and 48 transversions. The corrections were validated by Sanger sequencing at representative sites (Additional File 1: Fig. S6A, Additional File 7: Table S6). Among these corrected sites, 1,696 were located in genic regions, with 645 in exons and 1,051 in introns (Figure 5A). Errors in gene exons could lead to inaccuracies in the predicted CDS (Figure 5F). Using the polished genome, we re-predicted CDS for 645 genes with errors in their exons, resulting in 438 genes acquiring more accurate and extended CDS.

To update the functional annotation, predicted proteins were blasted against multiple public protein databases (Figure 1D, Additional File 1: Fig. S7A). In total, we annotated 25,846 functional genes, featuring an additional 1,732 functional genes compared to TGD2021 (Additional File 1: Fig. S7A, B). In the case of these newly annotated genes, protein functional annotation revealed their distribution across distinct structural domain families, with a higher prevalence observed in certain families, such as the leucine-rich repeat domain, the cyclic nucleotide-binding domain, and the WD40/YVTN repeat-like-containing domain (Additional File 1: Fig. S7C). Three newly annotated proteins were associated with epigenetic regulation (Additional File 1: Fig. S7D). Two featured a histone H3 K4-specific methyltransferase SET domain homologous to MLL5 (KMT2E) that are critical for gene transcription regulation, cell cycle regulation (G1/S transition), and myoblast differentiation [56–59]. Another exhibited homology to the 16S rRNA m5C methyltransferase NSUN4, characterized by the presence of a RsmB domain [60].

In this version (draft v5) (Additional File 1: Fig. S1), we optimized TSS annotation for 3,937 genes with multiple eTSSs, manually re-annotated 1,752 duplicated genes and 15 universally alternatively spliced genes, and re-predicted CDS for 438 genes.

### Annotate transcript isoforms generated by alternative splicing

It has been reported that 1,286 *Tetrahymena* genes generate alternative splicing (AS) isoforms [26], but this information was not integrated into previous gene models and TGD2021 contained only 459 AS genes. With the gene model being highly optimized in this study, we identified all six types of alternative splicing, namely exon skipping, alternative last exon, intron retention, mutually exclusive exons, alternative 5’ splice site, and alternative 3’ splice site, in a total of 3,041 genes, generating 5,917 isoforms (Fig. 1E, 6B). Consistent with the previous report [26], intron retention was the dominant form of AS (Fig. 6B). The numbers of AS genes and isoforms in our annotation were much higher than those in TGD2021 (gene: 5,041 vs. 459, isoform: 5,917 vs. 516) (Fig. 6B). Of these, 876 genes exhibited no less than two AS isoforms. Each AS event was supported by DRS full-length reads spanning the intron-exon junctions (Fig. 6A). Gene loci representing each of the six different AS types were selected for RT-PCR analysis, validating the existence and structure of AS isoforms (Additional File 1: Fig. S8A-F, Additional File 8: Table S7).

**Figure 6.**
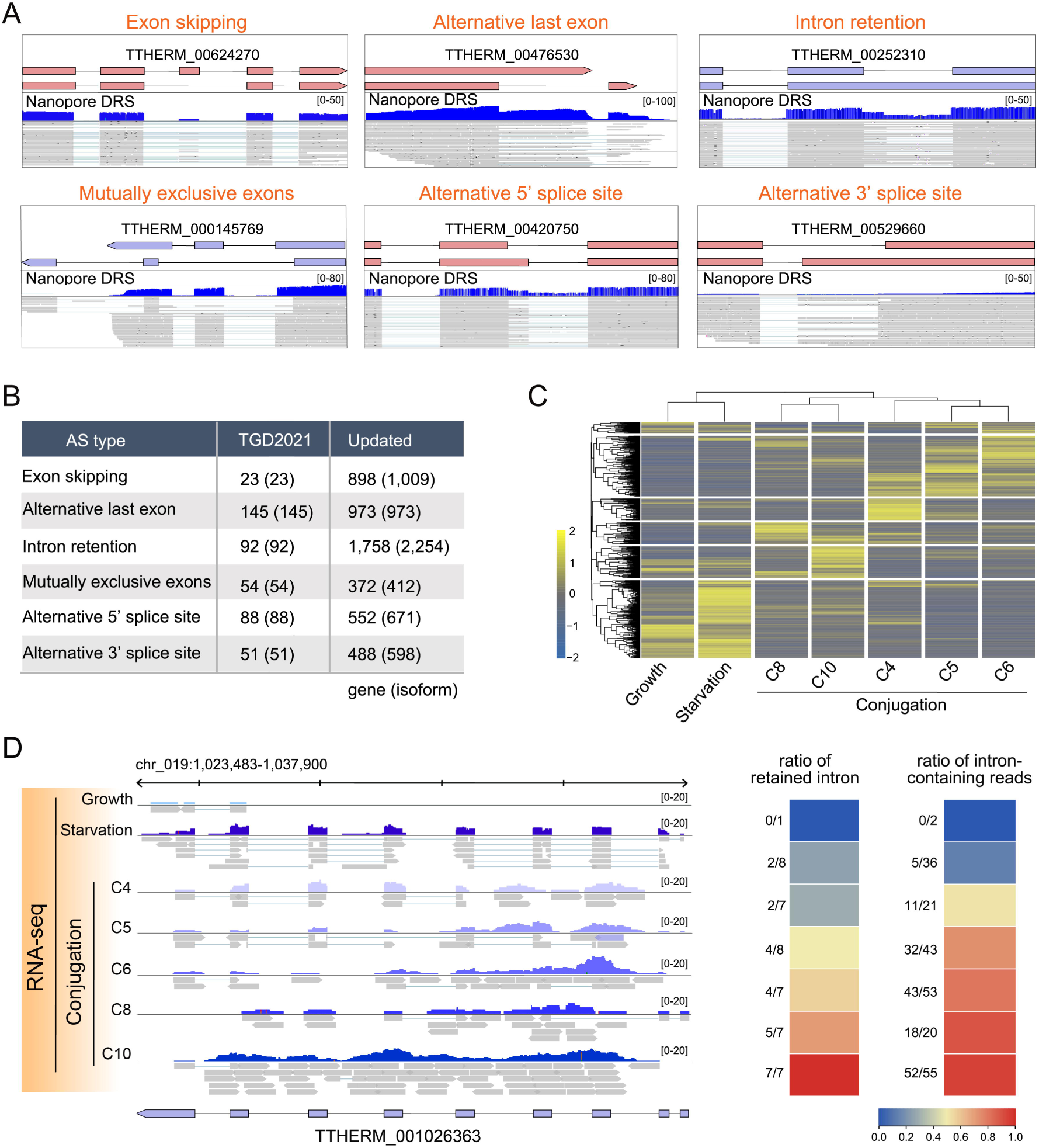
Annotation of alternatively spliced (AS) isoforms in *Tetrahymena*. A. The representative display of gene models and IGV snapshots of Nanopore DRS reads for six different AS types. B. Comparative summary of gene and isoform numbers in each of the six different AS types in TGD02021 and TGD2024 (updated). C. A heatmap depicting the expression profiles of AS transcripts across different stages: growth, starvation for 24h, and conjugation at 4h, 5h, 6h, 8h, and 10h. D. A representative gene exhibiting a stage-specific tendency for intron retention, supported by transcriptomic data (left), as well as the ratio of retained intron and the ratio of the intron-containing reads (right). The ratio of retained intron was defined as the retained intron number divided by the total sequenced intron number of the gene. The ratio of the intron-containing reads was defined as the reads aligned to the intron divided by the total reads aligned to the gene.

To further investigate whether the generation of AS isoforms was stage-specific, we compared AS isoforms in growth, starvation, and different timepoints of conjugation. The results showed that 2,131 out of 5,917 AS isoforms were generated across all periods, while others exhibited a tendency to be highly expressed during specific stages. Specifically, 114 AS isoforms were generated exclusively during growth, 326 during starvation, and 1,146 during conjugation (Fig. 6C). For example, in the case of the starvation- and conjugation-specific gene TTHERM_001026363, its AS isoforms showed a stage-specific pattern, with a gradual increase of the ratio of retained introns and intron-containing reads observed as conjugation progressed (Fig. 6D). In support, GO analysis of the overall functions of AS isoforms revealed a predominant enrichment in processes related to cell cycle and meiosis (Additional File 1: Fig. S8G, Additional File 9: Table S8).

### Identify natural antisense transcripts (NATs)

We observed that many gene loci could be transcribed from both sense and antisense strands (Fig. 7D-F). Intriguingly, transcripts originating from the antisense strand, which were typically shorter in length, were located within or in close proximity to the sense-coding transcripts, characteristic of natural antisense transcripts (NATs) [61]. In total, 4,389 NATs were identified (Fig. 7A). The presence of DRS and RNA-seq reads provided strong support for these NATs, confirming that they were *bona fide* transcripts rather than transcriptional noise. 14% of protein-coding genes (3,908/27,494) showed evidence of antisense transcription. Most NATs lacked a discernable ORF (>100 aa), but 11 NATs were annotated as potential functional protein and 112 displayed high protein-coding potentials (Additional File 1: Fig. S9A).

**Figure 7.**
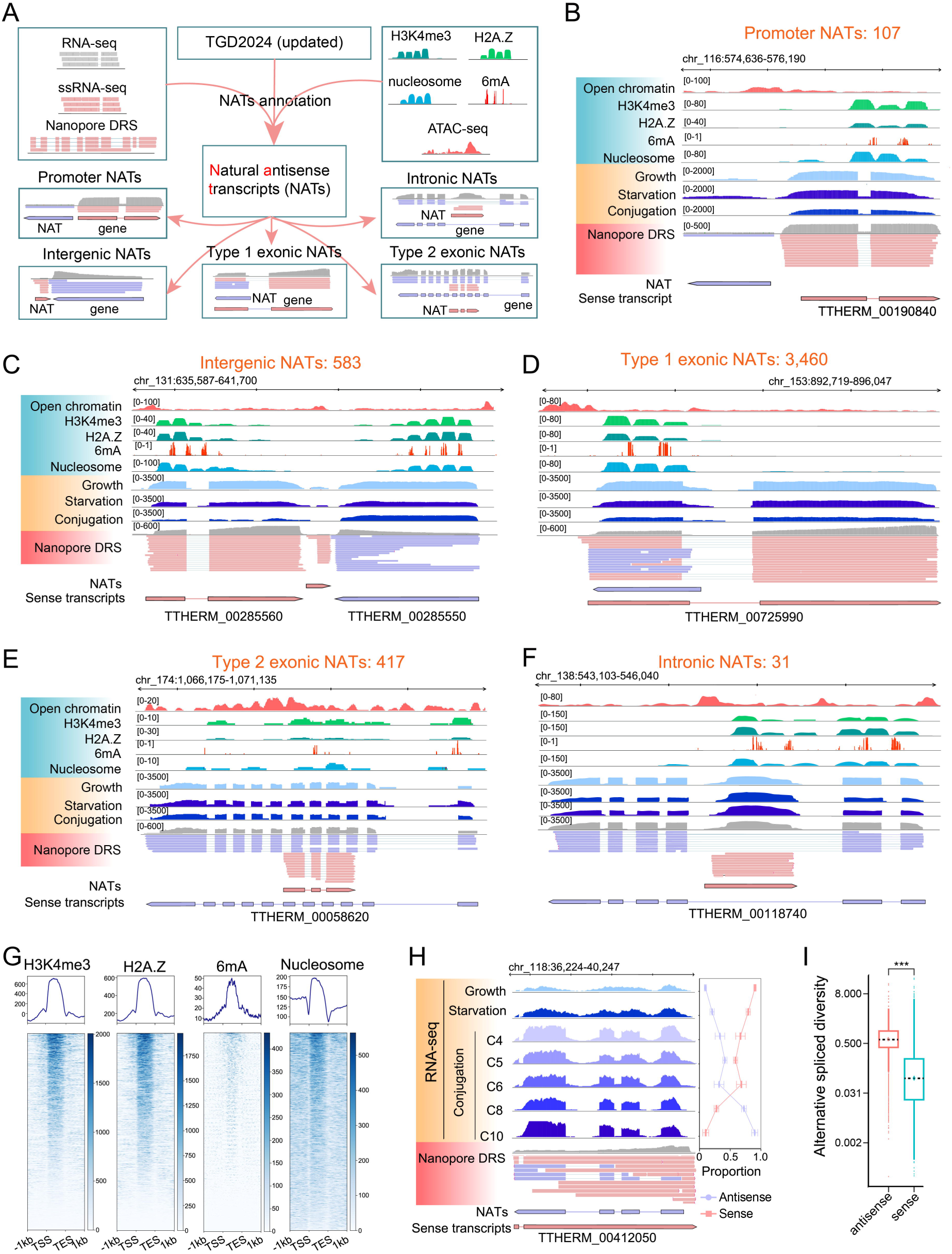
Identification and characterization of five types of natural antisense transcripts (NATs) in *Tetrahymena*. A. Schematics for NATs annotation. NATs were identified on the updated gene model (TGD2024) using both transcriptomic and epigenetic data. Identified NATs were further categorized based on their relative positions to corresponding sense transcripts. B-F. IGV snapshots showing five types of NATs. They included promoter NATs (B), originating from shared bidirectional promoters of the sense transcripts; intergenic NATs (C), transcribed from the upstream or downstream of the sense transcripts and possessing their own promoters; type 1 exonic NATs (D), located within 1 kb downstream of the TSSs of the sense transcripts and sharing epigenetic marks with their sense transcripts; type 2 exonic NATs (E), located more than 1 kb downstream of the TSSs of the sense transcripts; and intronic NATs (F), transcribed from the intronic regions of sense transcripts. G. Distribution profiles of H3K4me3, H2A.Z, 6mA, and well-positioned nucleosomes on the transcript body of NATs. Transcripts were scaled to unit length and was extended to each side by 1kb length. H. An IGV snapshot showing the anti-correlation of temporal expression patterns between a NAT and its corresponding sense transcript (left). The line chart (right) depicted the proportion of expression level for sense and antisense transcripts at different time points. The error bar represented the standard deviation (SD). I. The box plot showing that the alternative splicing diversity (ASD) of NATs exceeded that of their sense transcripts (the median of NATs and sense transcripts were 0.96 and 0.28, respectively). Student’s *t*-test was performed. *** *P* <0.001. ASD was defined as the number of different types of splice sites divided by the total reads aligned to the NATs or sense transcripts.

The resulting set of NATs was categorized into three groups according to their positional relationship with the corresponding sense transcript (protein-coding genes). 1) Intergenic NATs. These NATs did not overlap with other sense transcripts and were further subdivided into: a) 107 promoter NATs originated from shared bidirectional promoters (Fig. 7B), and b) 583 intergenic NATs possessing their own independent promoters (Fig. 7C). 2) Exonic NATs. These NATs were transcribed from loci with sense transcripts and were categorized into: a) 3,460 type 1 exonic NATs located within 1 kb downstream of the TSS of the respective sense transcription unit and shared the epigenetic marks with the latter (Fig. 7D), and b) 417 type 2 exonic NATs located more than 1 kb downstream of the TSS of the sense transcript and possessed the epigenetic marks downstream of their own TSS independently (Fig. 7E). It is worth mentioning that these NATs are not the reverse transcription or replication byproducts of sense transcripts. Instead, their exon-intron boundaries slightly deviate from those of their sense transcripts and they themselves contain canonical GU-AG sites for intron splicing (Additional File 1: Fig. S9B, C). 3) Intronic NATs. There were 31 NATs transcribed within the intronic regions of sense transcripts (Fig. 7F). The distinct genomic locations of NATs and their positional proximity to the sense transcripts may determine their roles in various aspects of gene expression.

Compared to sense transcripts, NATs were generally characterized by shorter lengths (Additional File 1: Fig. S9D) and lower expression levels (Additional File 1: Fig. S9E), while no difference in GC content was observed (Additional File 1: Fig. S9D). Interestingly, the majority of antisense transcripts also carried epigenetic marks (H3K4me3, H2A.Z, 6mA and well-positioned nucleosome) in their transcription units (Figure 7G, Additional File 1: Fig. S9F), similar to the report in *Arabidopsis thaliana* wherein NATs were enriched with H3K4me3 [62]. Most of these epigenetic marks were shared with their corresponding sense transcripts (Fig. 7D), but they were also possibly involved in regulating NATs expression.

Most importantly, these NATs exhibited temporal-specific expression patterns that were opposite to their corresponding sense coding genes, mirroring the findings in *Arabidopsis thaliana* where sense and antisense transcripts exhibited mutual exclusivity at individual loci [63]. In the instance of the gene TTHERM_00412050 in *Tetrahymena*, the expression of its NATs gradually decreased while the expression of its sense transcripts increased, during the transition from growth to starvation and subsequently to conjugation (Fig. 7H). This phenomenon might induce gene silencing of the corresponding sense genes by degrading the sense mRNA or interfering with its translation, a role that has been reported for NATs in plants [62, 64].

We then assessed the alternative splicing diversity (ASD), defined as the proportion of different types of introns for each NAT loci and its sense gene in DRS data. Intriguingly, the ASD of NATs significantly exceeded that of their sense counterparts (0.96 vs. 0.28, *P* <0.001) (Fig. 7I) or total sense coding transcripts (0.96 vs. 0.15, *P* <0.001) (Additional File 1: Fig. S9G). We speculate that, akin to non-coding RNA, NATs appear to be exempt from the evolutionary constraints imposed on protein-coding genes by the preservation of a functional ORF. This exemption might allow their exons to function as modular sections and act as independent units that can be shuffled and rearranged with great flexibility [55].

## CONCLUSION

In this study, we established a novel workflow to optimize the genome annotation of *Tetrahymena* that integrated large scale transcriptomic data, including RNA-seq data from different cell stages, strand-specific RNA-seq data, and Nanopore DRS data. Most importantly, epigenetic data including H3K4me3, H2A.Z, 6mA, and nucleosome along with ATAC-seq data were incorporated. This comprehensive dataset enabled the optimization of gene models, the accurate identification of TSS and TES, the augmentation of UTR information, the updated annotation of protein functions, and the addition of alternatively spliced isoforms.

Our updated gene model (TGD2024) (Additional File 1: Fig. S1) comprised a total of 27,494 genes, including 26,223 well-annotated and 1,271 low-confidence genes. Compared to TGD2021, we annotated 2,481 new genes, and optimized 6,428 genes including 3,878 exon-altered genes, 169 orientation-reversed genes, 65 TSS-altered genes, 1,379 fused genes, and 1,136 partitioned genes. We also increased the number of alternatively spliced isoforms to a total of 5,917 and annotated an additional 1,732 functional genes. Furthermore, we identified a large pool of NATs that might generate a diverse and extensive repertoire of potential regulatory RNAs.

Our work will largely facilitate *Tetrahymena* biology studies and the conceptual framework employed here holds substantial promise for facilitating genome annotation in other eukaryotes. Moving forward, we aim to enrich our analysis by incorporating additional epigenic data such as H3K27me1 that is predominantly enriched at the 3’ end of gene bodies (unpublished data) and Nanopore DRS data from conjugating cells. We also plan to generate data using QTI-seq and Cap-seq that will help to better determine the transcription start sites and translation start sites [65, 66].

## MATERIALS AND METHODS

### Cell growth, RNA extraction, and library construction

*Tetrahymena* wild-type strains (SB210 and CU428) were obtained from the *Tetrahymena* Stock Center (http://tetrahymena.vet.cornell.edu). Cells were grown in SPP medium at 30°C. For conjugation, starved SB210 and CU428 cells were resuspended in 10 mM Tris (pH 7.5) at 2 × 10^5^ cells/ml, mixed in equal volumes, and samples were collected at 4h, 5h, 6h, 8h, 10h after mixing. Total RNA was collected using the RNeasy Plus Mini kit (Qiagen, 74134). The quality and concentration of RNA samples were analyzed by 1% agarose gel electrophoresis and Qubit^®^3.0 Fluorometer (Thermo Fisher Scientific). Strand-specific RNA sequencing libraries and Illumina sequencing libraries of SB210 were constructed according to manufacturer-recommended protocols and sequenced by Novogene Co. Ltd (Beijing, China). For ATAC-seq libraries, transposase Tn5 from the Nextera DNA Library Preparation Kit (#FC-121-1030, Illumina, USA) was employed to treat 10^5^ macronuclei for 1h at 37°C. The DNA from the samples was subsequently recovered using the MinElute Recovery Kit (#28004, Qiagen, Germany). Amplification and library construction of the sample DNA were performed for 13 PCR cycles, with library adapter primers sourced from the Nextera XT Index Kit (#FC-121-1011, Illumina, USA). The DNA from the constructed library was recovered once more using the MinElute Recovery Kit (#28004, Qiagen, Germany).

### Gene loci identification based on RNA-seq and strand-specific RNA-seq data

The latest published MAC genome of *Tetrahymena thermophila* downloaded from TGD (http://ciliate.org) was used as the reference for reads mapping.

Adapters and low-quality reads were removed using Trim Galore (http://www.bioinformat-ics.babraham.ac.uk/projects/trim_galore/). Paired-end reads generated by RNA-seq and strand-specific RNA-seq were mapped back to the genome for transcript assembly using Hisat2 [67] with default parameters (-- rna-strandness R/RF for strand-specific RNA-seq). Picard Tools (https://broadinstitute.github.io/picard/) were used to remove duplicate reads from PCR. RNA-seq and strand-specific RNA-seq data were divided into three groups (growth, conjugation, starvation) according to the different cell cycle stages of *Tetrahymena*. The RNA-seq reads in each group were assembled and merged into the stage-combined transcripts by Stringtie with default parameters [68]. For each gene locus, only transcripts with the highest expression level (FPKM>1) across three groups were used for gene model optimization.

### Gene prediction and UTRs annotation using LoReAn2

The latest published MAC genome of *Tetrahymena* downloaded from TGD (http://ciliate.org) was used as the reference genome for reads mapping [24]. Gene prediction was performed using the LoReAn2 annotation pipeline [31]. In detail, the transcriptomic data were aligned to the genome using the Program to Assemble Spliced Alignments (PASA) [69] and the Genomic Mapping and Alignment Program (GMAP) [70]. For protein alignment, the analysis and annotation tool (AAT) [71] was used to align protein sequence to the genome. Reference genome-guided transcripts were assembled using Trinity [72]. SNAP [73], Augustus [74], and GeneMark-ET [75] were used to generate *de novo* gene annotation individually, which were then combined using EVM [69]. PASA was used to annotate UTRs.

### Reverse transcription polymerase chain reaction (RT-PCR)

Total RNA after Dnase treatment (Invitrogen, AM1907) was reverse-transcribed using an oligo-dT primer and M-MLV Reverse Transcriptase (Invitrogen, 28025013) and cDNA was used as a template. RT-PCR was performed using Premix Taq (TaKaRa, RR901A). All PCR primers are listed in Supplementary Data.

### ChIP-seq, MNase-seq, SMRT-seq data processing

The preprocessing steps for the epigenetic data (H3K4me3 ChIP-seq, H2A.Z ChIP-seq, MNase-seq) were consistent with those employed for RNA-seq. Only the mono-nucleosome sized fragments (120-260 bp) were analyzed. The unique mapping results (bam files) and the 6mApT sites (SMRT-seq 6mA data) were calculated by custom Perl scripts and plotted by deepTools [76] (bin = 10 bp).

### ATAC-seq and data processing

ATAC-seq was performed as previously described [77]. Libraries were sequenced (PE150) on an Illumina HiSeq sequencer. Mapped reads without PCR duplicates were used to retrieve short open chromatin regions (shorter than 100 bp) using deepTools [76], as defined by the Greenleaf research team [78, 79]. Peak calling was performed using MACS2 (v2.1.0) [80] The open chromatin profile distribution around TSS was plotted by deepTools [76]. The human precise TSSs from the RefTSS database [81] were mapped to corresponding genes using custom Perl scripts.

### Gene model optimization by a machine learning approach based on epigenetic information

To optimize the gene model predicted by transcriptomic data, a Random Forest (RF) model was developed to further identify TSS based on epigenetic information (H3K4me3, H2A.Z, 6mA, and well-positioned nucleosome). RF classification algorithm was implemented with the randomForest R package [82]. The model training was performed using a dataset containing abundant information of epigenetic marks in regions of 1,000 bp downstream of TSS (positive training set) and in regions of 1,000 bp centered by TES (negative training set). 10,460 well-annotated and longer than 1kb genes were selected, 70% of which were used for model training and the rest were for testing.

The error rate of the RF model was computed based on the out-of-bag (OOB) error, which is the mean prediction error over all Random Forest trees. The importance of each feature was computed as “mean decrease in accuracy” (MDA). Feature importance (MDA) and classification performance (OOB error) measures were further averaged over a collection of five hundred Random Forests to obtain stable results. The genome-wide regions were clustered and divided into different categories regions based on comprehensive consideration of chromatin states (H3K4me3, H2A.Z, 6mA, and nucleosome) using the DBSCAN cluster algorithm in fpc R package (eps=150, MinPts=3) (https://www.unibo.it/sitoweb/christian.hennig/en/). Subsequently, epigenetic mark signals on regions from clustered categories (scaled to 1kb) was feature engineered to predict TSS-regions using the pre-trained RF model.

### CDS prediction, UTR annotation and protein function annotation

Longest ORFs on each strand were predicted from the stage-combined transcripts using ORFfinder [83]. A putative ORF was defined as amino acid sequences exceeding 100 aa in length. The orientation of strand-specific RNA-seq transcripts was used to determine CDSs from longest predicted ORFs. The regions beyond CDSs on transcripts were defined as 5’ UTRs and 3’ UTRs, respectively. Putative protein coding regions were annotated using EggNOG [84], Interproscan [85], and Pannzer2 [86] by mapping to known proteins, protein domains and signal peptides collected in UniProtKB database [87], Pfam databases [88], and InterPro database [89].

### Motif enrichment analysis

For the motif enrichment analysis of open chromatin region upstream of the TSS, only fragments shorter than the mono-nucleosome size (100 bp) were analyzed after mapping to the genome. Peaks were identified using MACS2 (v2.1.0) [80]. Motif enrichment analysis was performed in called peaks using HOMER (v4.8) [90].

For the motif analysis around the TES, sequences spanning 50 bp upstream and downstream of RNA cleavage sites/TES were extracted using bedtools [91]. Subsequently, the extracted fasta sequences were renamed utilizing SeqKit [92]. Motif analysis was conducted on these sequences using MEME-Suite’s simple enrichment analysis (SEA) [93].

### Nanopore direct RNA sequencing data generation and analysis

Oxford PromethION 2D amplicon libraries for full-length transcriptome sequencing were generated according to the Nanopore community protocol using library preparation kit SQK-LSK109 and were sequenced on R9 flowcells to generate fast5 files. Fastq files were derived from fast5 reads by basecalling using guppy v3.2.10 (default parameters, https://github.com/metagenomics/denbi-nanopore-training/blob/master/docs/basecalling/basecalling.rst). Reads were filtered using Nanofilt v2.5.0 [94]. Nanopore direct RNA sequencing (DRS) reads were aligned to the genome using minimap2 v2.16 [95]. The alternatively spliced isoforms were identified by customized Perl scripts followed by manual curation. The RNA-seq data (growth, starvation 24h, and conjugation at 4h, 5h, 6h, 8h, and 10h), each with two or three biological replicates, were used to calculate the expression level during different cell stages. The heatmap plot and gene ontology (GO) analysis were plotted by Tbtools [96]. For the poly-A tail analysis, nanopolish-polya version 0.10.2 (https://github.com/jts/nanopolish) was used to estimate polyadenylated tail lengths from Nanopore DRS raw reads.

### Identification and classification of natural antisense transcripts

Natural antisense transcripts (NATs) were identified by fulfilling the following criteria: 1) transcribed from the antisense strand of protein-coding genes as evidenced by DRS data, and 2) localized upstream or within protein-coding genes, encompassing intronic or exonic regions. Classification of each transcript as either coding or noncoding was determined using a stepwise filtering pipeline. First, all candidates were scored with LGC [97] to determine their coding potential. All transcripts that were named “non-coding” were retained as potential noncoding candidates. Second, all candidate transcripts were subjected to blastp [98] and HMMER (versus Pfam-A and Pfam-B) [99]. For blastp and HMMER, transcripts were translated in all three sense frames. Transcripts with an E-value less than 1e-4 in any of the three search algorithms were considered as functional-coding; transcripts that were predicted to contain ORF exceeding 200 bp in length, yet lacked identifiable homologous proteins or functional domain, were defined as potential-coding; and the remaining were classified as non-coding. The alternative splicing diversity (ASD) was quantified as the ratio between the number of distinct splice sites and the total reads number captured by Nanopore DRS data for a particular gene. The comparison of ASD for sense protein-coding genes and NATs could be approached in two ways: 1) comparing all NATs (4,398) to protein-coding genes with NATs (4,398); and 2) comparing all NATs (4,398) to all protein-coding genes (27,494).

## Supplementary information

**Additional file 1: Figure S1.** The flowchart for genome annotation. **Figure S2.** RT-PCR validation of gene models optimized by the transcriptomic data. **Figure S3.** The eTSSs/pTSSs prediction and gene optimization using epigenetic data. **Figure S4.** RT-PCR validation of gene models optimized by eTSSs. **Figure S5.** The regulatory elements of untranslated regions analysis in *Tetrahymena*. **Figure S6.** Four types of error sites polished by Illumina and Sanger sequencing data. **Figure S7.** Protein function annotation on draft v4. **Figure S8.** The experimental validation and functional analysis for alternative splicing (AS) isoforms. **Figure S9.** Identification and characterization of natural antisense transcripts (NATs).

**Additional file 2: Table S1**. The Information of the sequencing data.

**Additional file 3: Table S2**. The comparison between TGD2014, TGD2021, and the LoReAn2 annotated gene model.

**Additional file 4: Table S3**. PCR primers for gene optimization.

**Additional file 5: Table S4**. Homology protein of the promoter binding protein in *Tetrahymena*.

**Additional file 6: Table S5**. Results of GO enrichment analysis for gene with poly-A.

**Additional file 7: Table S6**. PCR primers for genome polish.

**Additional file 8: Table S7**. PCR primers for alternative splicing.

**Additional file 9: Table S8**. Results of GO enrichment analysis for alternative splicing.

## Data availability

Scripts to generate data and to perform the above analysis are available in github: https://github.com/yefei521/UTR-annotation. Public RNA-seq datasets were available on the Gene Expression Omnibus (GEO) under accession number GSE27971: https://www.ncbi.nlm.nih.gov/gds/?term=GSE27971 [26]. Our SMRT-seq 6mA data was available at the NCBI database (BioProject accession number: PRJNA932808) [30], MNase-seq nucleosome data was available at https://www.ncbi.nlm.nih.gov/sra/SRX5146438[accn] [28]. ChIP-seq data (H3K4me3 and H2A.Z), RNA-seq, strand-specific RNA-seq, Nanopore direct RNA sequencing, and ATAC-seq data from the current work were deposited at the NCBI database (BioProject accession number: PRJNA1048844).

## Ethics approval and consent to participate

Not applicable.

## Consent for publication

Not applicable.

## Author contribution

Fei Ye conceived and led the project, performed strand-specific RNA-seq and Nanopore direct RNA sequencing, conducted computational and experimental analysis, and wrote and revised the manuscript. Xiao Chen provided analytical framework for TSS prediction and ATAC-seq processing, conducted ATAC-seq, revised the manuscript, and provided funding resources. Yalan Sheng assisted in resolving analytical issues and revised the manuscript. Lili Duan and Aili Ju performed RNA-seq of conjugation samples. Naomi Stover assisted with gene nomenclature and formatted data for presentation on TGD. Khaled A. S. Al-Rasheid revised the manuscript. Shan Gao conceived and supervised the project, wrote and revised the manuscript, and provided funding resources. All authors read and approved the final manuscript.

## Acknowledgements

We thank members of the Gao laboratory for discussion. Our special thanks are given to Dr. Weibo Song (Ocean University of China, OUC) for his kind suggestions during preparation of the manuscript and Dr. Jun Wang (Institute of Microbiology, Chinese Academy of Sciences) for his advice on Nanopore direct RNA sequencing and analysis. We appreciate the computing resources provided by the Institute of Evolution & Marine Biodiversity in OUC, the Center for High Performance Computing and System Simulation at Laoshan Laboratory, and Marine Big Data Center of Institute for Advanced Ocean Study of OUC.

---

**Figure S1.**
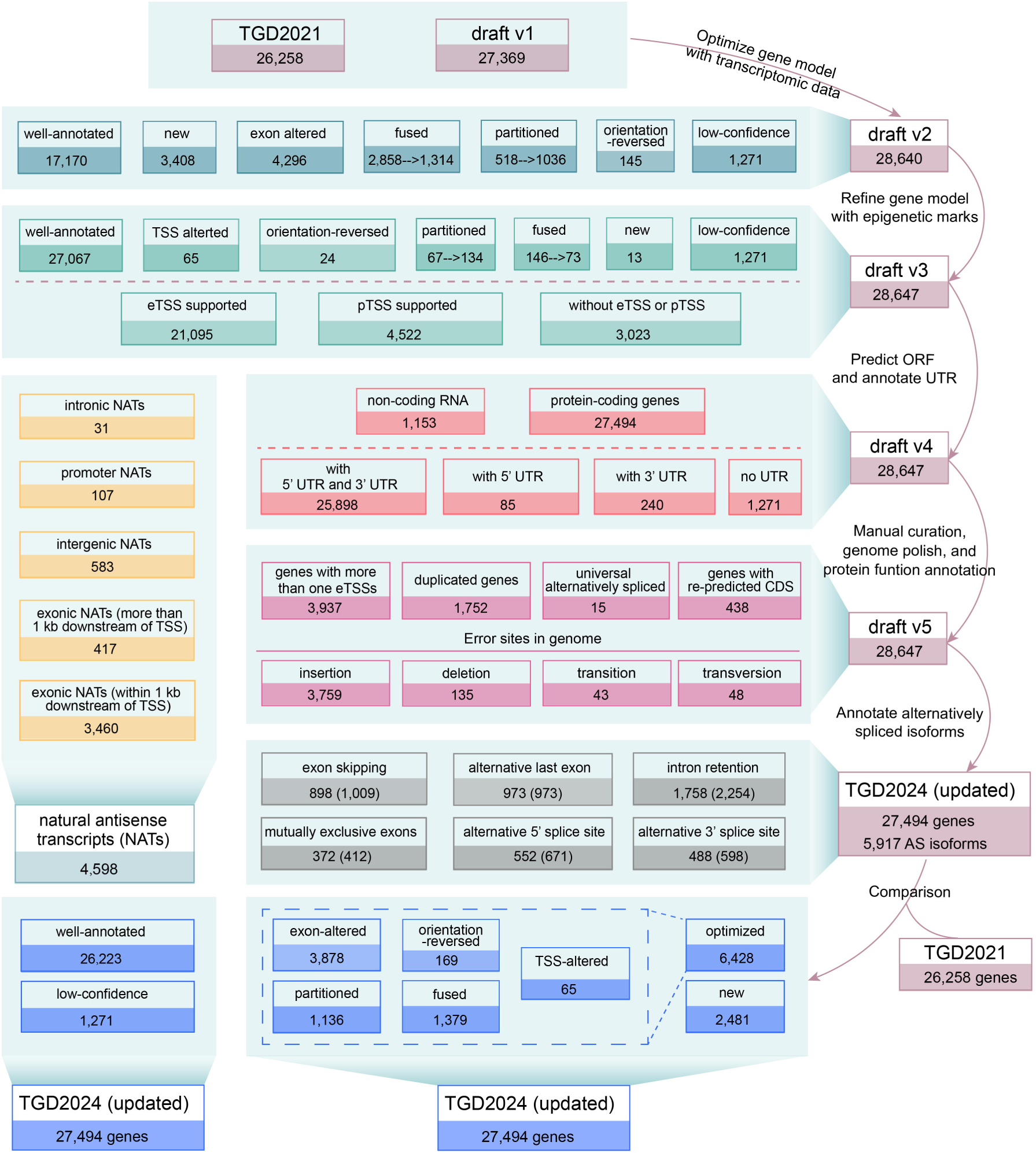
The flowchart for genome annotation.

**Figure S2.**
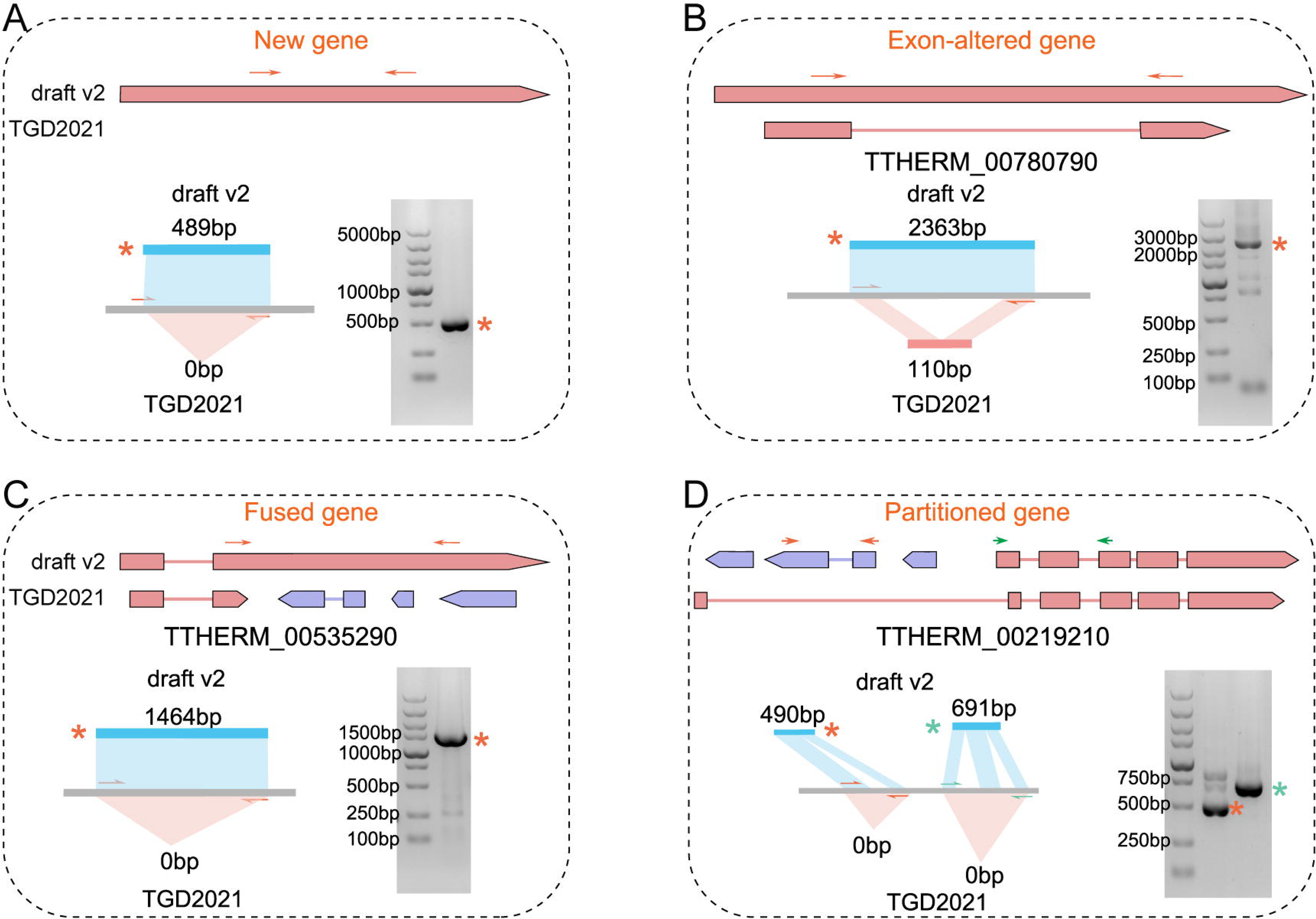
RT-PCR validation of gene models optimized by the transcriptomic data, including new genes (A), exon-altered genes (B), fused genes (C), and partitioned genes (D). For each representative gene, gene model in draft v2 and TGD2021 (top), primers flanking the target region and the expected size of RT-PCR products (bottom left), and the gel electrophoresis image of RT-PCR products (bottom right) were displayed. Orange and green stars indicated corresponding RT-PCR products.

**Figure S3.**
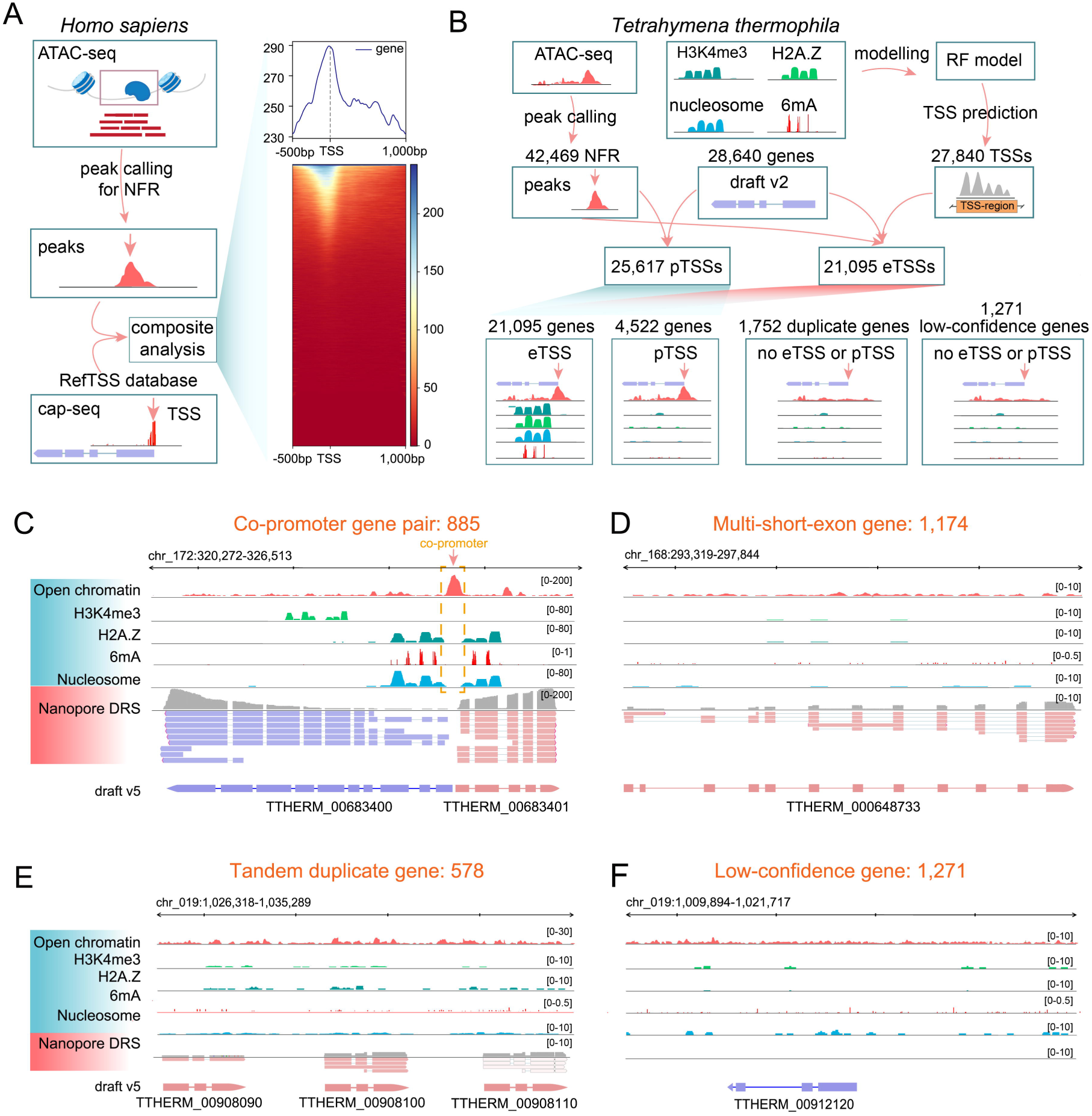
The eTSSs/pTSSs prediction and gene optimization using epigenetic data. A. Peaks corresponding to nucleosome free regions (NFRs) were identified using *Homo sapiens* ATAC-seq data and were subsequently integrated with TSS data from RefTSS for analysis. The results indicated that ATAC-seq fragments from NFRs tended to be enriched around TSS. B. The prediction of eTSSs and pTSSs using epigenetic data. Genes were classified into different groups according to the presence or absence of eTSSs and pTSSs. C-F. IGV snapshots of different types of genes optimized by epigenetic data, complemented by transcriptomic data, including gene pairs sharing their promoters (C), genes with multiple short exons (D), and tandem duplicate genes (E). Low-confidence genes (F) were neither supported by epigenetic marks nor Nanopore DRS reads.

**Figure S4.**
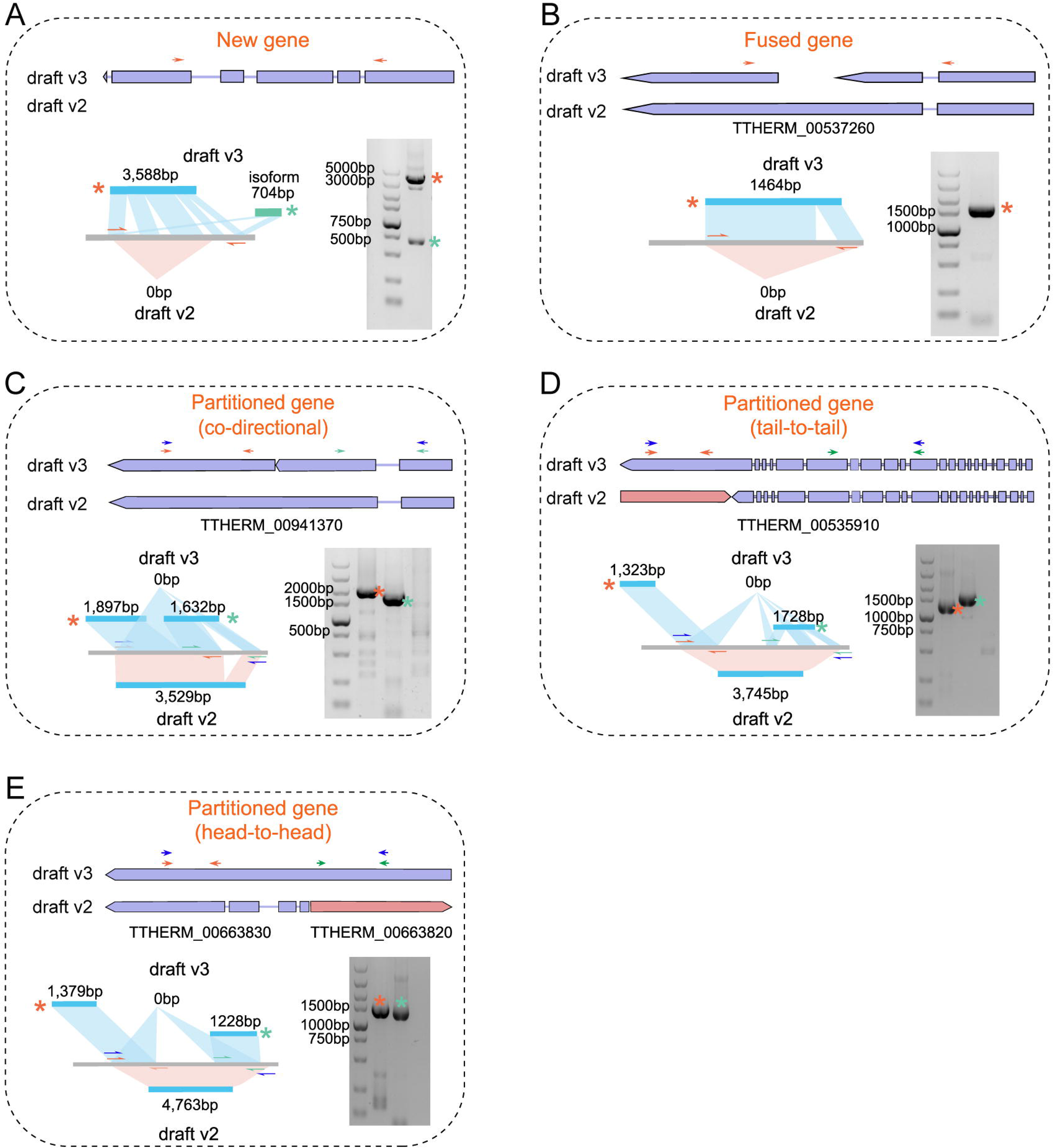
RT-PCR validation of gene models optimized by eTSSs, including new genes (A), fused genes (B), and co-directional partitioned genes (C), tail-to-tail partitioned genes (D), and head-to-head partitioned genes (E). For each representative gene, gene models in draft v2 and draft v3 (top), primers flanking the target region and the expected size of RT-PCR products (bottom left), and the gel electrophoresis image of RT-PCR products (bottom right) were displayed. Orange and green stars indicated corresponding RT-PCR products.

**Figure S5.**
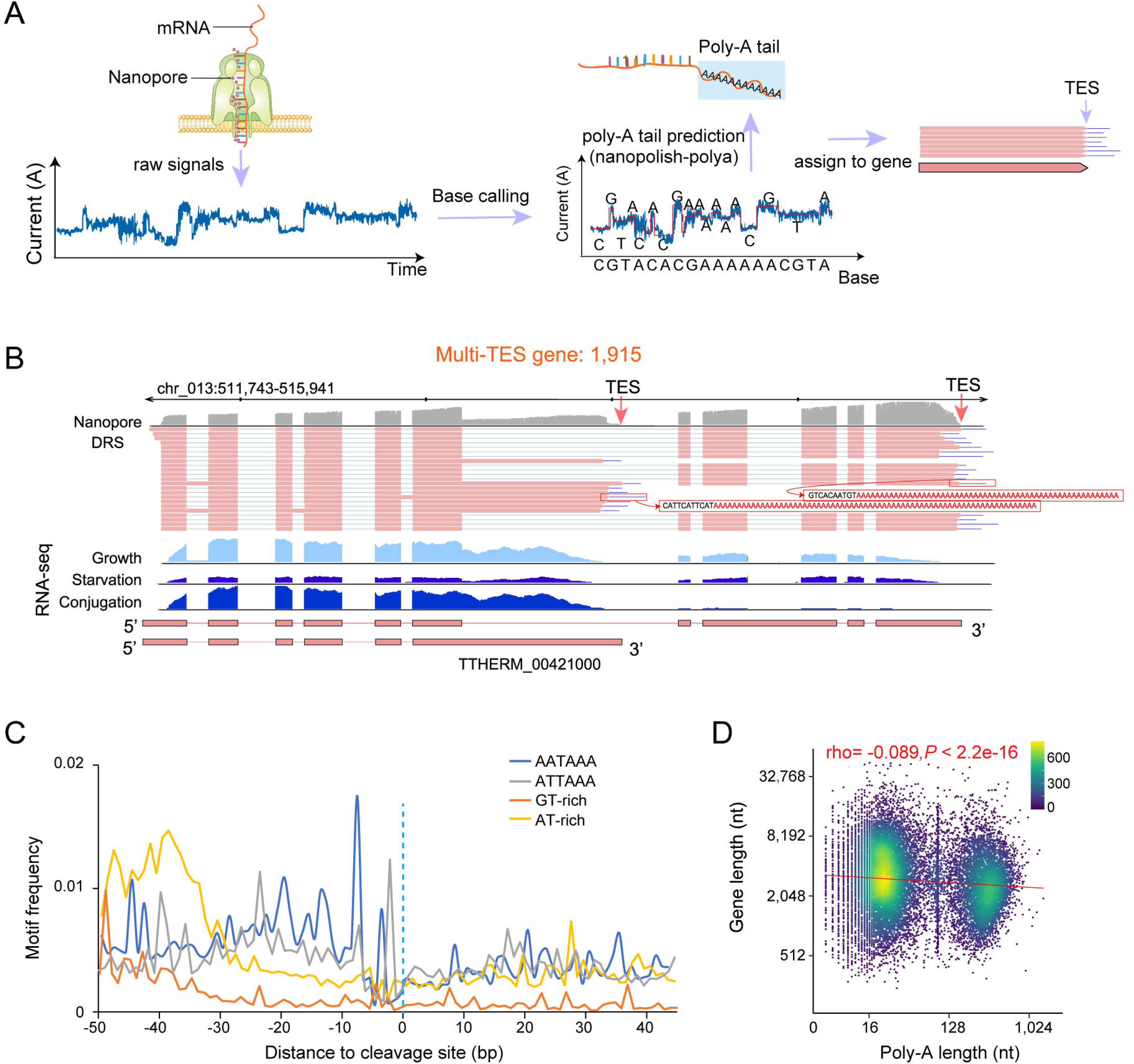
The regulatory elements of untranslated regions analysis in *Tetrahymena*. A. The workflow of Nanopore sequencing and the prediction of poly-A tails. B. An IGV snapshot of a gene with multiple TESs, identified according to the position of poly-A tails in DRS reads. C. Distribution frequency of main regulatory motifs around cleavage sites. The dominant motifs of PASs (AATAAA and ATTAAA) were enriched within 20 bp upstream of cleavage sites. The regulatory elements such as GTGT (GT-rich elements) and TGTA (AT-rich elements) were enriched more than 30 bp upstream of cleavage sites. D. The Spearman’s correlation showed no correlation between poly-A tail length and gene length (rho= -0.089, *P* <2.2e-16). The longest poly-A tail was selected as the representative for each gene. Both axes were plotted on a logarithmic scale.

**Figure S6.**
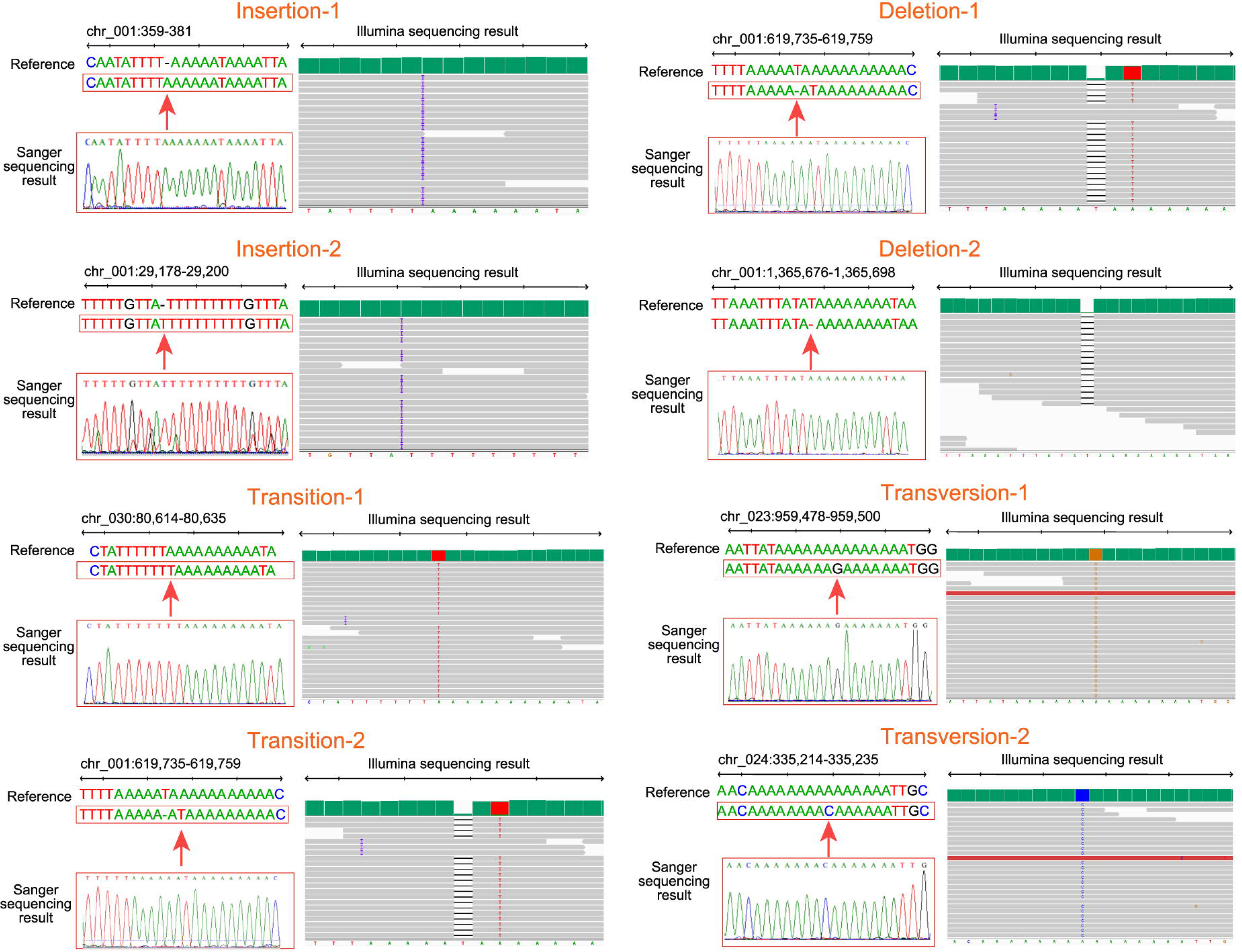
Four types of error sites polished by Illumina and Sanger sequencing data, including two examples each for insertion, deletion, transition, and transversion.

**Figure S7.**
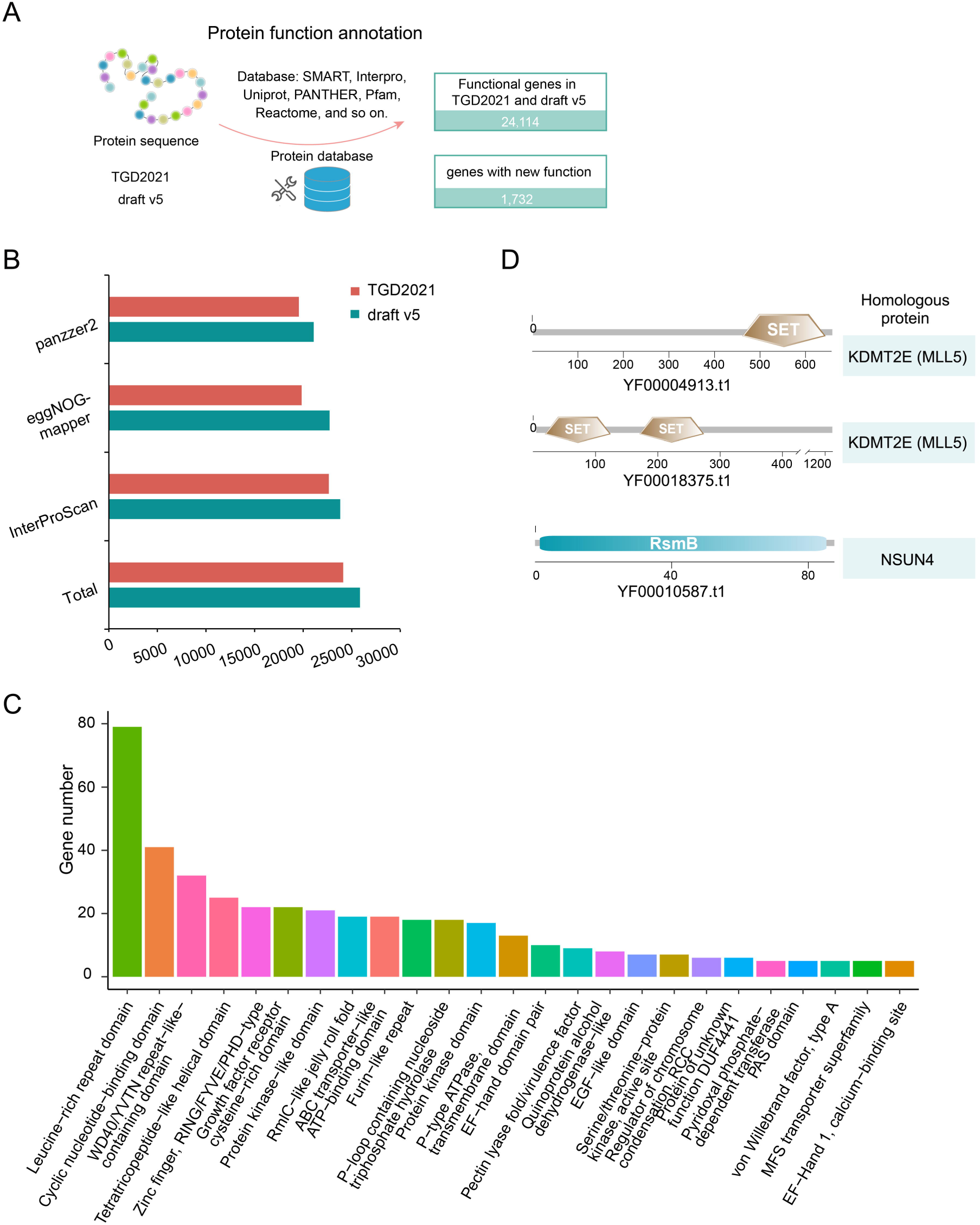
Protein function annotation on draft v4. A. Protein sequences of TGD2021 and draft v4 were blasted against multiple public protein databases, featuring an additional 1,732 functional genes compared to TGD2021. B. The number of functional genes annotated by different databases for TGD2021 and draft v5. C. Top 25 functional gene groups for genes with new functions. D. Domain structure of four newly annotated proteins possibly related to epigenetic regulations, homologous to MLL5 and NSUN4, respectively.

**Figure S8.**
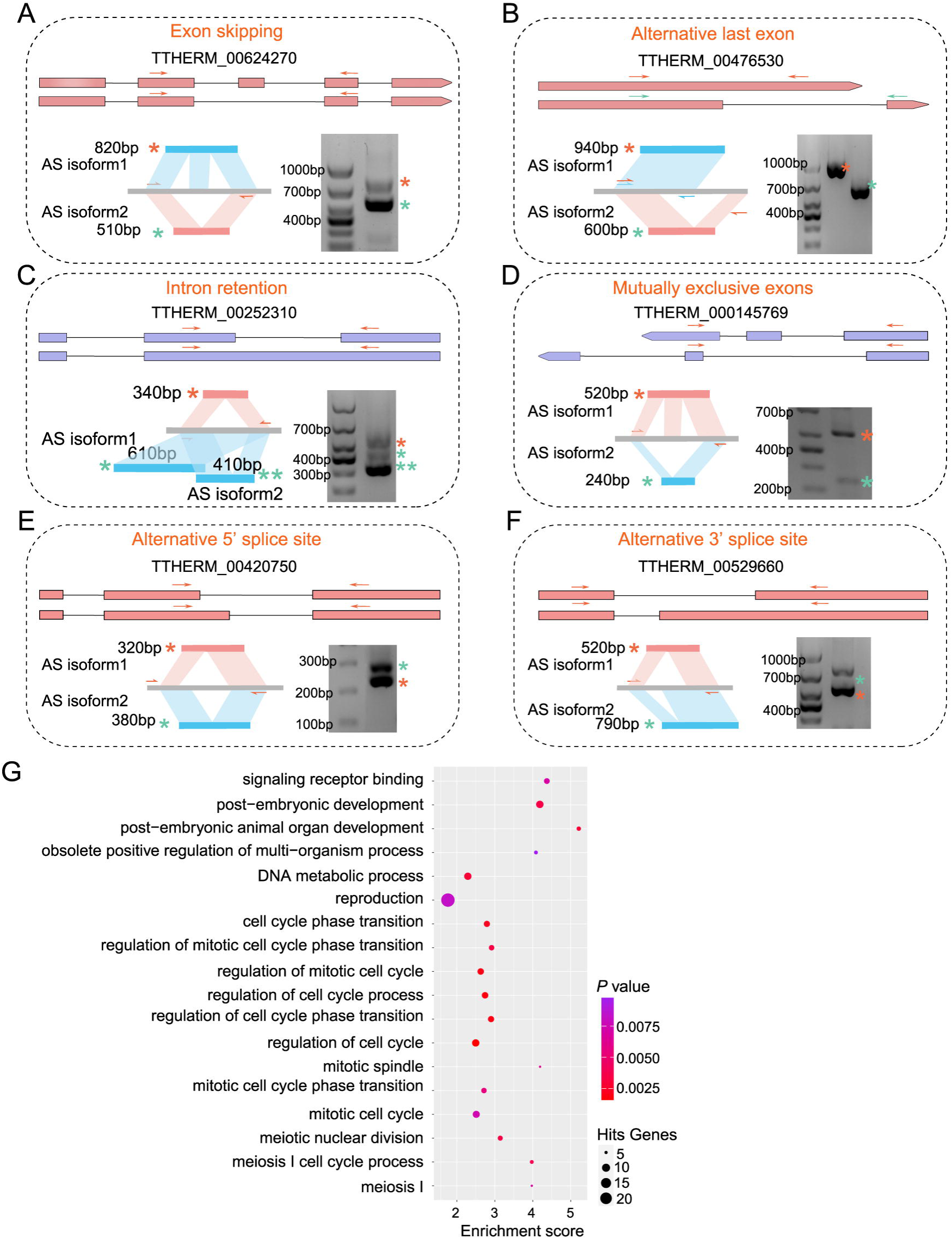
The experimental validation and functional analysis for alternative splicing (AS) isoforms. A-F. RT-PCR validation of the presence of AS transcripts, including exon skipping (A), alternative last exon (B), intron retention (C), mutually exclusive exons (D), alternative 5’ splice site (E), and alternative 3’ splice site (F). For each representative gene, gene model in TGD2024 (updated) and isoform, primers flanking the target region and the expected size of RT-PCR products (bottom left), and the gel electrophoresis image of RT-PCR products (bottom right) were displayed. Orange and green stars indicated corresponding RT-PCR products. G. GO analysis of total AS isoforms revealed that genes possessing AS isoforms were primarily enriched in processes related to cell cycle regulation.

**Figure S9.**
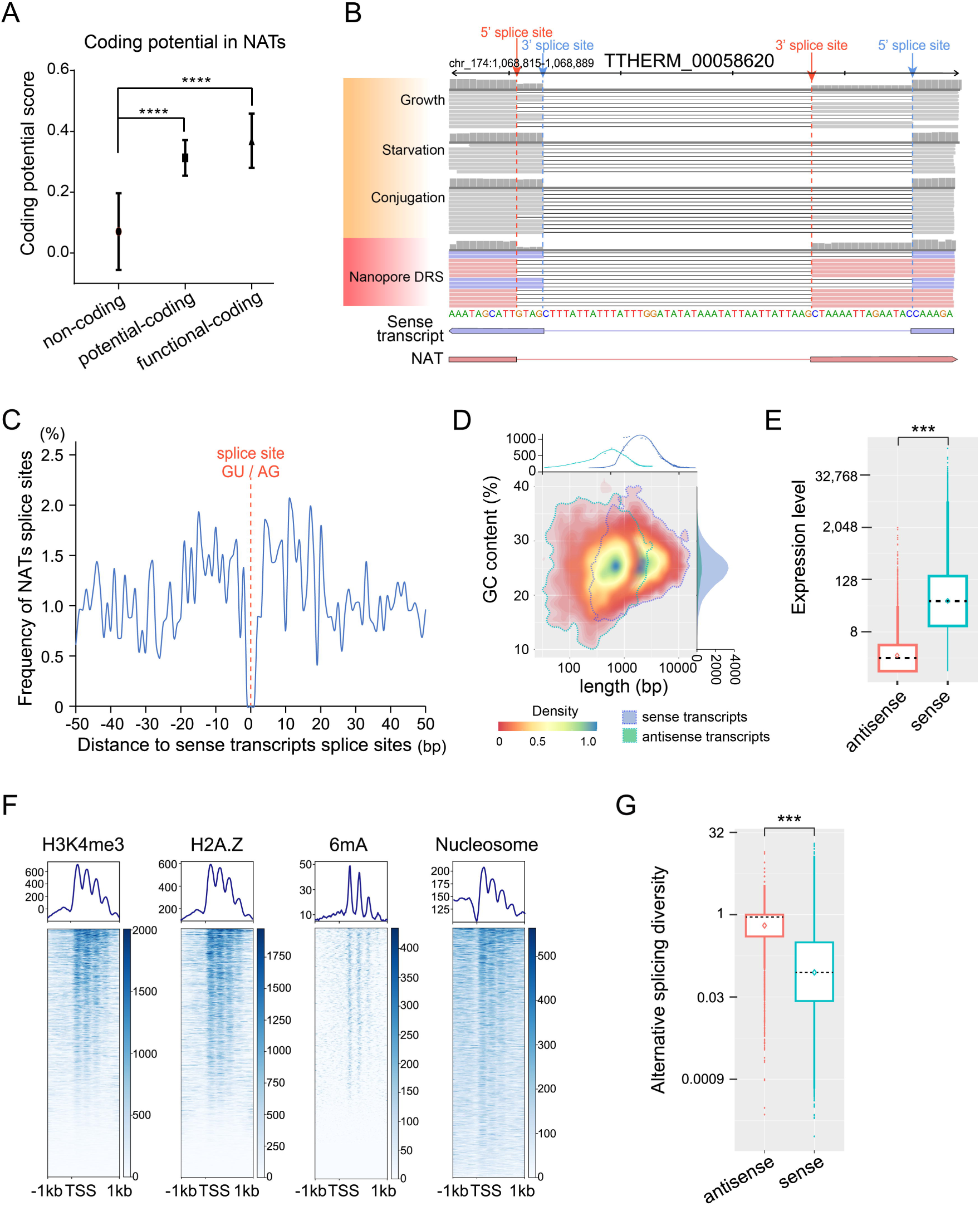
Identification and characterization of natural antisense transcripts (NATs). A. NATs were divided into non-coding, potential-coding and functional-coding, according to their significantly different coding potential scores. ****, *P*<0.0001. B. An IGV snapshot showing the splice sites of a NAT (red) and its corresponding sense transcript (purple). Note that this NAT possessed its own canonical GU-AG site for intron splicing and its exon-intron boundary slightly deviated from that of its sense transcript. C. Distribution of intron-exon boundaries of NATs in relation to those of sense transcripts. The red dashed line at position 0 marked the location of the sense intron-exon boundaries. D. The heatmap showing the GC content and length of sense transcripts and antisense transcripts. NATs had shorter length but had no difference in GC content comparing to sense transcripts. E. The box plot showing that NATs had lower expression level comparing to sense transcripts. The expression level was defined as DRS reads counts of NATs and its corresponding sense transcripts. ***, *P*<0.001. F. Composite distribution profiles of H3K4me3, H2A.Z, 6mA, and nucleosome near the TSS of NATs. The distribution profiles showing that all four marks were accumulated downstream of TSS. G. The box plot showing that the alternative splicing diversity (ASD) of total NATs exceeded the total sense transcripts (the median of NATs and sense transcripts were 0.96 and 0.15, respectively). Student’s *t*-test was performed. ***, *P* <0.001. ASD was defined as the number of different splice sites divided by the total reads aligned to the NATs or sense transcripts.

